# Tabular Foundation Models Are Competitive Cellular Perturbation Predictors Across Biological Scales

**DOI:** 10.64898/2026.06.28.735106

**Authors:** Giovanni Palla, Alexander Hillsley, Yang-Joon Kim, Loic A. Royer

## Abstract

Predicting how cells respond to genetic and chemical perturbations is a central challenge in drug discovery and functional genomics. A growing ecosystem of specialized single-cell foundation models has been developed to address this problem, yet their practical advantage over domain-agnostic approaches remains unclear. Here we evaluate the power of Tabular Foundation Models such as TabICL and TabPFN, general-purpose pre-trained regression models, against domain-specific architectures including PRESAGE, scGPT, scLAMBDA, STACK and Prophet across four complementary evaluation settings: cell-level in-context cross-cell-type prediction, pseudobulk perturbation prediction on five Perturb-seq datasets of cell-lines, a genome-wide CRISPR screen in primary human CD4^+^ T cells, and embryo-level cell-type composition prediction in a zebrafish developmental perturbation atlas. In the cell-level cross-cell type perturbation prediction, Tabular Foundation Models perform on par or better than specialized models. On pseudobulk perturbation prediction, Tabular Foundation Models consistently out-perform specialized baselines across multiple evaluation metrics and datasets. On whole-embryo cell-type composition prediction, Tabular Foundation Models are competitive with specialized baselines. These results demonstrate that general-purpose tabular in-context learning provides a strong and scalable alternative to bespoke biological architectures for perturbation response modeling across cell systems and scales.

## 1 Introduction

Predicting how cells respond to genetic perturbations is a central goal of functional genomics and a prerequisite for rational drug design. High-throughput CRISPR screens coupled with single-cell transcriptomics (Perturb-seq) now routinely profile thousands of perturbations in a single experiment [Replogle et al., 2022, Nadig et al., 2025, Zhu et al., 2025, Saunders et al., 2023]. Similar multiplexing scale have been achieved for chemical or biological perturbations [Zhang et al., 2025, Anchang et al., 2024].

The prevailing approach to perturbation prediction has been to design specialized architectures with explicit biological inductive biases [Lotfollahi et al., 2023, Cui et al., 2024, Bunne et al., 2023, Klein et al., 2025, Palla et al., 2025, Adduri et al., 2025, Dong et al., 2026, Littman et al., 2025, Ji et al., 2025]. These models typically aim to predict the gene expression response to a perturbation either in a new cellular context (e.g., a new cell type) or for a new, unseen perturbation. The former approach leverages in-context information from related cells in the target system, while the latter relies on transferable gene- or perturbation-level representations – derived from prior knowledge – as encoders of perturbation effects.

Despite these architectural differences, all of these methods can be viewed within a common regression framework. Let *c* denote the control cellular context, *p* the perturbation, and *y* the prediction target. Each model aims to estimate the conditional mean response

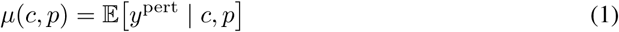

The definition of *y* depends on the biological scale of interest: for single-cell models such as STACK, *y* = *x* is the single-cell gene expression profile; for pseudobulk models such as PRESAGE, *y* = *x̄* is the pseudobulk expression profile; and for phenotype-level models such as Prophet, *y* = *s* is a higher-level functional signature, such as a cell-type composition vector or functional assays readouts. Under this view, the main differences between these approaches lie in their inductive biases and representations of *c* and *p*, rather than in the statistical object they seek to predict. Across these settings, models in the field have focused on learning the conditional mean response using various inductive biases and representations of *c* and *p*.

This perspective also suggests why tabular foundation models should be strong candidates for perturbation prediction. Prior-Fitted Networks (PFNs) [Müller et al., 2024] perform approximate Bayesian posterior predictive inference at test time: given a training dataset D, a single forward pass approximates

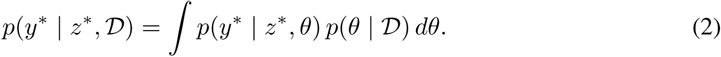

The corresponding posterior predictive mean, 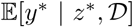, is precisely the quantity required for perturbation prediction. This is especially appealing in our setting, where the number of observed perturbations is limited, the covariates are naturally heterogeneous and tabular, and test perturbations are often encoded with a diverse set of representations. In such regimes, strong posterior predictive regression can matter more than hand-crafted biological architecture. Tabular foundation models [Hollmann et al., 2025, Qu et al., 2026] therefore offer a compelling alternative: they amortize inference over a broad family of regression problems, can be highly data-efficient, and do not require biology-specific pretraining to achieve competitive predictions.

In this work, we show that *Tabular Foundation Models* (TFMs), general-purpose pretrained regression models trained on synthetic data, match or outperform all tested specialized architectures for perturbation prediction across cell systems and biological scales. We evaluate TFMs in four settings: cell-level perturbation prediction for chemical perturbations, pseudobulk perturbation prediction on five Perturb-seq cell-line datasets, a genome-wide CRISPR screen in primary human CD4^+^ T cells, and embryo-level cell type composition prediction in a zebrafish developmental perturbation atlas. Across all four settings, TFMs consistently match or outperform every specialized baseline we tested. These results suggest that biology-specific inductive biases are not required to obtain strong perturbation predictors, and that a single pretrained tabular model, combined with standard dimensionality reduction, can serve as a competitive and scalable alternative to domain-specific models. Code is available at https://github.com/royerlab/tfm-perturbation.

## 2 Related Work

### Perturbation prediction models

Computational approaches for predicting cellular responses to perturbations have progressed from linear and additive models [Ahlmann-Eltze et al., 2025, **?**] to deep generative and foundation-model-based methods [Bunne et al., 2023, Theodoris et al., 2023, Cui et al., 2024]. For example STACK [Dong et al., 2026] uses in-context learning with tabular attention across cellular contexts, metagene tokenization, and a negative binomial likelihood to model single-cell responses. PRESAGE [Littman et al., 2025] learns a conditional mapping from perturbation embeddings – derived from gene interaction networks, biological foundation models, and literature embeddings – to perturbed expression profiles. Concurrent work [Cole et al., 2026] has further shown that prior-knowledge-based biological foundation model embeddings, combined with attention-based fusion, can approach the experimental noise floor on some benchmarks. scGPT [Cui et al., 2024] and scLAMBDA [Wang et al., 2024] pretrain transformers on millions of single-cell profiles using expression tokenization schemes and then fine-tune them for perturbation prediction. Prophet [Ji et al., 2025] combines multiple sources of evidence, together with transcriptomic data, to predict higher-level functional readouts under genetic and chemical perturbations. Taken together, different models are designed for the predictive task they are applied to, and indeed entail specialized architectures and pretraining.

### Tabular foundation models and prior-fitted networks

In parallel, a separate line of work has developed foundation models for generic tabular prediction tasks. Prior-fitted networks (PFNs) [Müller et al., 2024] introduced the idea that a transformer trained on synthetic datasets sampled from a prior over supervised learning problems can perform approximate Bayesian posterior predictive inference at test time, without gradient-based adaptation on the target task. This view reframes prediction as amortized inference over a family of regression problems rather than optimization on a single dataset. TabPFN citepHollmann2025 showed that this strategy can yield strong performance on small tabular problems, especially in low-data regimes where classical model selection is difficult. Subsequent work argued that prior-fitted prediction represents a broader paradigm for Bayesian inference [Müller et al., 2025], while TuneTables [Feuer et al., 2024] improved scalability through context optimization. More recent studies have incorporated causal structure [Swelam et al., 2025] and causal data augmentation [Bühler et al., 2026] to improve robustness, and TabICLv2 [Qu et al., 2026] currently represents the state of the art open model for tabular regression. These developments are directly relevant to perturbation prediction, where training perturbations are often limited, covariates are heterogeneous, and held-out interventions can be structurally novel. In such settings, a model that is strong at posterior predictive regression on structured tabular data is a natural candidate to be competitive even without biology-specific pretraining.

## 3 Background

### 3.1 Perturbation prediction as regression

Perturbation prediction can be formulated at multiple biological scales (Figure 1). At the *cell level*, given drug-treated source cells and untreated target cells of a different type, the goal is to predict what drug-treated target cells look like. At the *pseudobulk level*, given aggregated treatment effects for training perturbations, the goal is to predict the treatment effect for held-out perturbations. At the *organism level*, the target may be a non-expression readout such as cell type composition or other functional assays.

**Figure 1:**
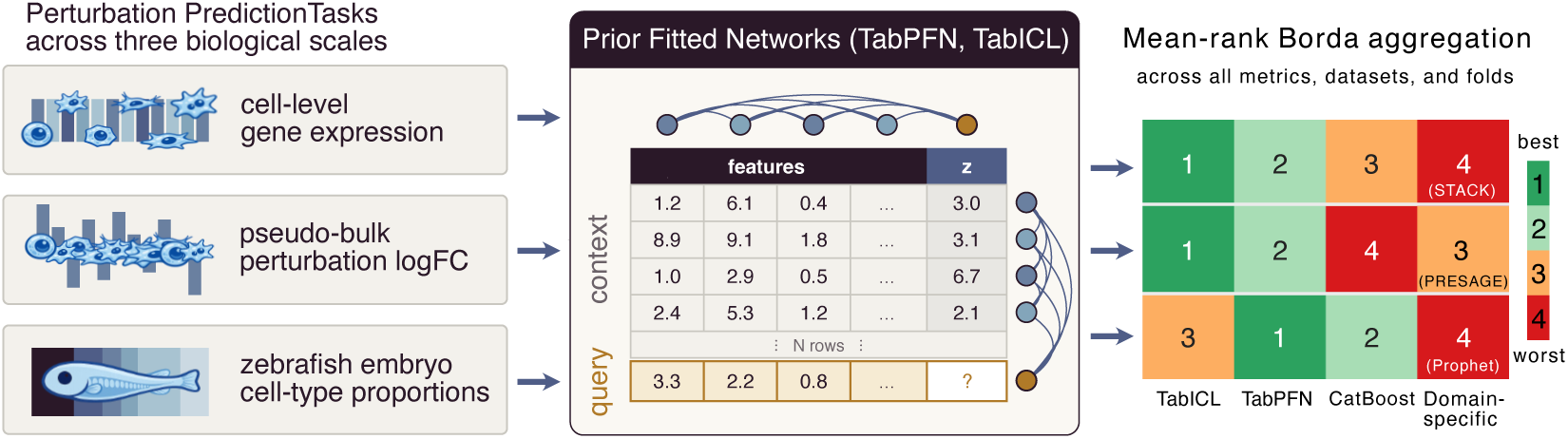
Unified Tabular Foundation Model (TFM) framework for perturbation prediction across three biological scales. **Left:** three task settings, each defining a different notion of “row” — OT-matched cell pair (cell-level gene expression), perturbation (Perturb-seq pseudo-bulk logFC), and perturbation × timepoint (zebrafish embryo cell-type proportions). **Centre:** all settings are funnelled into the same tabular regression substrate. The frozen TabPFN/TabICL backbone interleaves 1D feature attention (across columns) and 1D sample attention (across rows) followed by a per-cell MLP, and predicts the query row’s scalar target *z* from a context of *N* training rows — one PCA component at a time. **Right:** per-task ranking obtained by mean-rank Borda aggregation across all metrics, datasets, and folds, comparing two PFN backbones (TabICL, TabPFN), a strong tabular baseline (CatBoost), and the domain-specific model for each task (STACK for cell-level, PRESAGE for pseudo-bulk, Prophet for zebrafish). Cells are colour-coded green (best, rank 1) to red (worst, rank 4); a PFN backbone ranks first on every task — TabICL on the cell-level and pseudo-bulk settings, TabPFN on zebrafish.

Specialized models (PRESAGE, scGPT, scLAMBDA, STACK, Prophet) address these tasks through domain-specific architectures that operate in the full gene space, jointly predicting all output dimensions through *multi-output regression*. Tabular foundation models (TFMs), by contrast, are *univariate regressors*: they predict a single scalar target from a feature vector and a labeled context set. Below we describe how we adapt TFMs to each setting.

### 3.2 Cell-level prediction via optimal-transport matching

In the cell-level setting, the goal is to predict single-cell expression profiles for a target cell type given drug-treated source cells and untreated target cells. TFMs serve as componentwise regressors, where each context row is an *individual cell*.

#### Cell matching

Let **S**_ctrl_ ∈ ℝ*^n_s_^*^×^*^G^* denote source (e.g., T cell) control cells and **T**_ctrl_ ∈ ℝ*^n_t_^*^×^*^G^* denote target (e.g., NK cell) control cells. We fit PCA on the pooled control cells to obtain a shared embedding space ℝ*^d^*. We then solve an optimal-transport problem between the source and target control distributions in PCA space, computing a transport plan ***π***^∗^ that pairs source cells with target cells based on transcriptomic similarity:

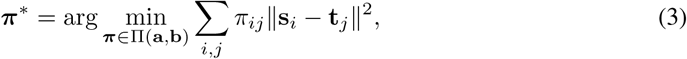

where Π(**a**, **b**) is the set of transport plans with uniform marginals. This yields *n* matched pairs {(**s***_i_,* **t***_σ_*_(*i*)_)}, where *σ* is the assignment derived from ***π***^∗^.

#### Componentwise cell-level regression

The matched control pairs define the training data for the TFM. Let **s***_i_*^pca^ ∈ ℝ*^d^* and **t***_σ_*_(*i*)_^pca^ ∈ ℝ*^d^* be the PCA projections of matched source and target control cells. For each PCA component *k*:

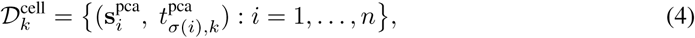

where each row is a single cell and the target is the *k*-th PCA component of the matched target cell. The TFM learns a mapping from source to target cell space using these pairs as context. At test time, drug-treated source cells {**s***_j_*^drug^} are projected to PCA space and passed through the TFM:

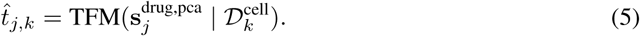

Repeating for all *k* and applying inverse PCA yields predicted drug-treated target cell profiles 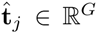. This produces a *distribution* of predicted cells (one per source drug cell), capturing cell-to-cell heterogeneity. The context set here contains ∼2,000–5,000 matched cell pairs.

#### Underlying assumption

This pipeline rests on the assumption that the cross-cell-type translation is approximately invariant to the perturbation: the source → target mapping learned from DMSO control pairs is reused at test time on drug-treated source cells. The assumption is reasonable when the perturbation-induced shift is small relative to the cross-cell-type shift, which is what it is usually observed in both genetic and chemical perturbations datasets. Interestingly, we do observe that the matching algorithm of choice has an impact in performance, where the OT match is superior to a knn based match (Table 10).

### 3.3 Pseudobulk prediction via componentwise regression

At the pseudobulk level, we predict perturbation-level treatment effects. For each perturbation *p*, aggregating cells yields ***δ****_p_* ∈ R*^G^* (perturbed mean minus control mean). We decompose the *G*-dimensional target into *d_y_* independent scalar regressions via PCA, where each context row is a *perturbation*.

#### Output decomposition

Let **Δ** ∈ ℝ*^K^*^train×^*^G^* be the matrix of training treatment effects, where *K*_train_ is the number of training perturbations and *G* the number of genes. We fit PCA to obtain **W***_y_* ∈ ℝ*^G^*^×^*^dy^* (*d_y_* ≪ *G*, typically 128) and project each perturbation’s response: **z***_p_* = **W***_y_*^⊤^***δ****_p_* ∈ ℝ*^dy^*.

#### Perturbation features

Each perturbation *p* is represented by a feature vector **x***_p_* ∈ ℝ*^dx^*, constructed from one or more embedding modalities. In the *multi-modality* setting, we reuse the same input modalities utilized by PRESAGE [Littman et al., 2025], which consists of gene-level representations from multiple sources (gene co-expression, protein interaction networks, pathway memberships, etc.) that are each independently PCA-projected and concatenated into a single feature vector per perturbation. In the *perturb-seq* settings features are derived from gene co-expression: we transpose the training treatment effect matrix to ℝ*^G^*^×^*^K^*^train^, apply PCA to obtain gene-level embeddings **E** ∈ ℝ*^G^*^×^*^dx^*, and look up the embedding of each perturbation’s target gene: **x***_p_* = **E***_g_*_(*p*)_, where *g*(*p*) is the gene targeted by perturbation *p*.

#### Componentwise in-context learning

For each output component *k* ∈ {1*, . . ., d_y_*}, we form a tabular dataset:

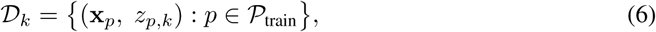

where **x***_p_* ∈ ℝ*^dx^* is the feature vector and *z_p,k_* ∈ ℝ is the *k*-th principal component of the treatment effect. Each row is a perturbation; the context set typically contains ∼50–500 rows. The TFM predicts the target for a query perturbation in a single forward pass:

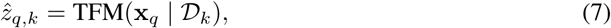

without any gradient update—the pretrained weights are frozen. This is repeated independently for each *k*, yielding 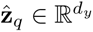, which is reconstructed via 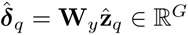.

The full pipeline thus makes *d_y_* TFM calls, each on a small tabular dataset (∼100 rows, ∼64 features, 1 target), the regime where prior-fitted networks excel [Hollmann et al., 2025, Qu et al., 2026].

#### Unifying observation

The TFM architecture and inference mechanism are identical across settings: in the cell-level benchmark each context row is a matched cell pair, in pseudobulk it is a perturbation, and in zebrafish it is a perturbation-timepoint pair predicting the corresponding cell-type composition. Only the feature representation and scalar response change across tasks.

#### Single-cell

We evaluate cell-level prediction on the OpenProblems cross-cell-type benchmark (298K cells from 3 donors, 144 drug conditions, 4 cell types), where models train on T cell responses and predict NK, B, and Myeloid responses on a 5k-HVG panel. We compare TabICL, TabPFN, and CatBoost under the same OT-matching pipeline against STACK, a single-cell foundation model with tabular attention [Dong et al., 2026]. Metrics are Pearson Δ, DE Spearman, direction match, DE recall, discrimination score, and Pearson E-distance (a Pearson correlation between per-perturbation real and predicted energy distances; see Appendix C); full preprocessing and evaluation details are given in Appendix A and Appendix C.

#### Pseudobulk

We evaluate perturbation-level prediction on five Perturb-seq datasets (HepG2, Jurkat, K562 Essential, K562 Genome-Wide, and RPE1 Essential) using replicate-aware 5-fold CV, and on a genome-wide CRISPR screen in primary CD4^+^ T cells from two donors [Zhu et al., 2025] as a primary-cell extension of the same pseudobulk setting. The CD4 benchmark uses within-donor perturbation holdout, with guide-level Stim48hr-versus-Rest log-fold-change vectors on the top-5k HVGs as targets and a multi-modal perturbation feature set. We compare TabICL, TabPFN, and CatBoost to PRESAGE, scGPT, scLAMBDA, and the Mean and Bootstrap controls, where Bootstrap is an oracle upper bound that resamples cells from the held-out perturbation’s own data (Appendix B.10). Metrics are relative MSE on the top-20 DE genes, cosine similarity, phenocopy AUROC, and Recall@10; dataset and metric details are in Appendix A and Appendix C.

#### Zebrafish

To test whether the same tabular formulation transfers beyond transcriptomic response prediction, we evaluate on the zscape developmental perturbation atlas [Saunders et al., 2023] : ∼2.7M cells, 28 genetic perturbations, 5 timepoints (18–72 hpf), and 98 cell types (after dropping rare cell type/timepoint pairs with ≤ 10 cells). The target is the perturbation- and timepoint-specific 98-dim cell type composition vector. Tabular regressors (CatBoost, TabICL, TabPFN) operate on a 398-d feature vector (intervention + phenotype embeddings) per (perturbation, timepoint) condition with PCA decomposition on both X and Y (*d_x_* = *d_y_* = 32), placing them on the same multi-output footing as Prophet. Evaluation uses 5-fold strict timepoint holdout (each fold removes all rows at one developmental timepoint). We compare TabICL, TabPFN, and CatBoost to Prophet [Ji et al., 2025] together with Mean baseline, and report Spearman correlation and *R*^2^ as in Appendix A.

#### Implementation

For all experiments we used the TabICLv2 [Qu et al., 2026] and TabPFN 2.6 [Hollmann et al., 2025] weights, without fine-tuning or test-time adaptation. Both TabICL and TabPFN are pretrained Prior-Fitted Networks [Müller et al., 2024]. This makes TFMs very competitive compared to specialized architecture, because they are explicitly designed to leverage the in-context information and do not require expensive post-training or fine-tuning.

## 4 Results

### 4.1 Cell-level cross-context perturbation prediction

We first evaluate cell-level perturbation prediction, where models are trained on perturbations from one context (cell type, cell line) and must predict perturbation responses in a held-out settings. Table 1 compares two PFN networks (TabICL and TabPFN), one strong tabular regressor (CatBoost) and one in context learning model designed for the task and pre trained on a large collection of scRNA-seq data (STACK). We used the OpenProblems 2k-HVG benchmark [Anchang et al., 2024] (see Appendix A.4 for the dataset and Appendix A.5 for the cell-matching pipeline). In order to apply PFNs to this task, we introduce an adaptation step to match single-cell gene expression profiles between resting cells of the two types under consideration. TabICL (OT), leads on Pearson Δ (0.570) and perturbation discrimination (0.795), TabPFN (OT) leads on DE Spearman (0.682), and CatBoost (OT) leads on DE recall (0.751); the three tabular regressors are within SEM on Pearson Δ and DE Spearman. STACK retains an edge on direction match (0.491) and Pearson E-distance (0.679).

**Table 1:**
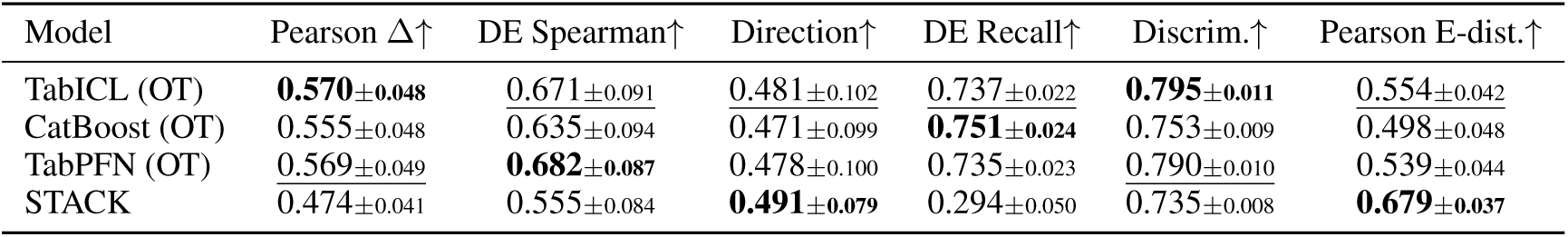
Cell-level cross-cell-type prediction on the OpenProblems 2k HVG benchmark. Mean ± SEM across 3 target cell types (NK, B, Myeloid) for Donor 1, with optimal-transport (OT) cell matching. Best per column in bold, second-best <u>underlined</u>.

| Model | Pearson $\Delta\uparrow$ | DE Spearman↑ | Direction↑ | DE Recall↑ | Discrim.↑ | Pearson E-dist.↑ |
| --- | --- | --- | --- | --- | --- | --- |
| TabICL (OT) | **0.570±0.048** | <u>0.671±0.091</u> | 0.481±0.102 | <u>0.737±0.022</u> | **0.795±0.011** | <u>0.554±0.042</u> |
| CatBoost (OT) | 0.555±0.048 | <u>0.635±0.094</u> | 0.471±0.099 | **0.751±0.024** | 0.753±0.009 | <u>0.498±0.048</u> |
| TabPFN (OT) | <u>0.569±0.049</u> | **0.682±0.087** | 0.478±0.100 | 0.735±0.023 | <u>0.790±0.010</u> | 0.539±0.044 |
| STACK | 0.474±0.041 | 0.555±0.084 | **0.491±0.079** | 0.294±0.050 | 0.735±0.008 | **0.679±0.037** |

OT matching consistently outperforms nearest-neighbour matching across both regressors (Appendix, Table 10): Pearson Δ gains of +0.07 (TabICL) and +0.11 (CatBoost). To assess robustness to context size, we sweep the fraction of source cells from 10% to 100% (Appendix, Figure 5). The OT regressors degrade gracefully, while STACK is more sensitive to context size. Overall these results indicate that TFMs are able to leverage the in-context information and, despite not being trained on the data, are superior to domain-specific knowledge trained on the same data modality.

### 4.2 Pseudobulk perturbation prediction

#### 4.2.1 Unseen perturbation prediction in cell-lines

We next evaluate PFNs on pseudobulk perturbation prediction across five Perturb-seq datasets using the benchmark protocol introduced by PRESAGE [Littman et al., 2025]. Results are reported in Figure 2. Each point in the summary figure represents a model’s mean performance across 5-fold CV on one dataset; per-dataset per-fold breakdowns are provided in Appendix Figure 6 and Table 8 (multimodal +Perturb-seq features) and Figure 7 and Table 9 (only multi-modal features). The Bootstrap baseline serves as an oracle upper bound, since it resamples cells from the held-out perturbation’s own data (see Appendix B.10; the resulting gap to the best learned model is tabulated in Table 6). Among learned models, TabICL and TabPFN consistently achieve the best performance, with TabICL leading on relative MSE and TabPFN performing comparably. scGPT performs near the Mean baseline on most datasets, while scLAMBDA shows intermediate performance, suggesting a potential inability to train on pseudobulked scRNA-seq data.

**Figure 2:**
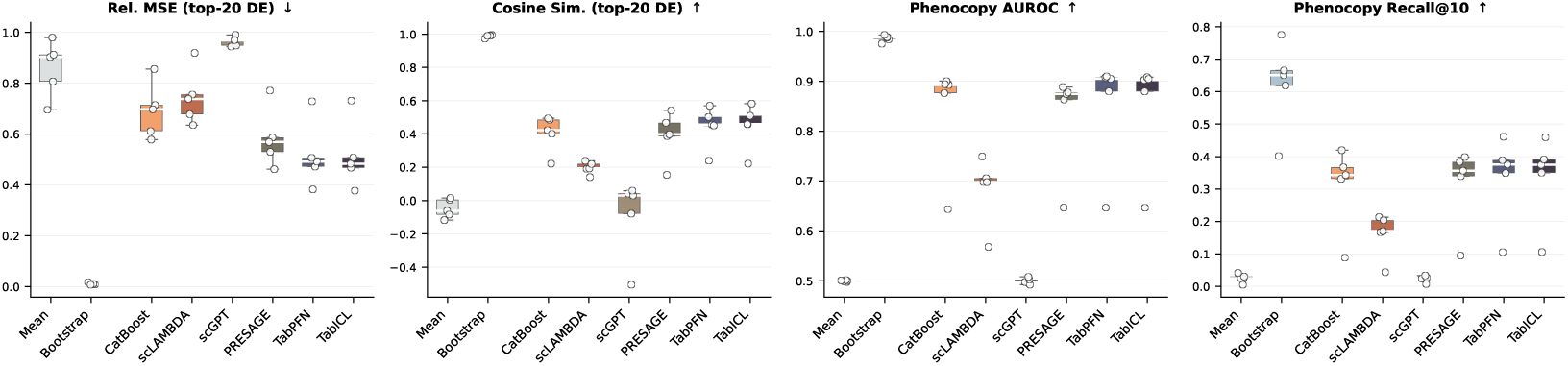
Pseudobulk perturbation prediction summary across five Perturb-seq datasets. Each point represents one dataset’s mean across 5-fold CV; boxplots show the distribution across datasets. The Bootstrap serves as an oracle upper bound (Appendix B.10). Among learned models, TabICL and TabPFN consistently outperform PRESAGE, scLAMBDA, scGPT, and CatBoost. Per-dataset per-fold results in Appendix Figures 6 and 7.

Concatenating Perturb-seq pseudobulk references from non-held-out cell lines on top of the multimodal embeddings improves both TabICL and PRESAGE performance (Appendix, Figure 7): for instance TabICL achieves 0.468 relMSE on HepG2 with the combined Perturb-seq + multi-modal features, versus 0.549 with multi-modal embeddings only - a 17% improvement on this dataset. The pattern is consistent across datasets and models and aligns with what was originally reported by PRESAGE [Littman et al., 2025]: same-modality (gene-expression) features from related cell lines carry the bulk of the perturbation-prediction signal, and PFNs effectively leverage it when concatenated with auxiliary multi-modal embeddings (Appendix D.4). A leave-one-out ablation across the six top embedding modalities (Appendix Figure 11) further shows that no single auxiliary modality dominates – individual drops produce changes within roughly ±1% relMSE in most cells.

We further isolate the contribution of the output-space PCA decomposition (Appendix G.2, Figure 9): replacing PCA with five controlled alternatives at matched rank *d_y_* = 128 shows that projecting onto bottom-128 PCs or a random 128-d Gaussian subspace collapses cosine-top-20-DE (Δ ≤ −0.31), a random orthogonal rotation of the PCA basis is nearly a no-op (Δ = +0.019), and non-orthogonal NMF unexpectedly outperforms PCA by +0.117 on average, hence what matters is that the output basis covers the high-variance directions of the target, not PCA *per se*. Performance is also robust to the precise PCA rank: a sweep over input (*d_x_*) and output (*d_y_*) dimensionality plateaus past *d_y_* ≈ 48 and is optimal around *d_x_* = 64 (Appendix Figure 10). Together with the support-set sensitivity, this points to the in-context inference step, rather than a reusable per-perturbation encoding, as the key aspect of the TFMs’ advantage.

Finally, we evaluate data efficiency by training each model on 25%, 50%, 75%, and 100% of the available training perturbations (Figure 8, Appendix). TabICL demonstrates strong performance even with limited training data-at 25% training data, it often matches or exceeds the performance of PRESAGE and CatBoost trained on the full dataset. PRESAGE shows the steepest performance gains with increasing data, consistent with its conditional framework requiring more examples to learn the perturbation manifold.

#### 4.2.2 Unseen perturbation prediction in Genome-wide CD4**^+^** T cell screen

We next evaluate on the genome-wide CD4^+^ T cell Perturb-seq screen from Zhu et al. [2025], which profiles CRISPR knockouts of ∼11k expressed genes in primary human T cells across two donors (D1, D2). Per donor, we restrict to perturbations whose knockout significantly downregulated their own target gene (≈ 7k of 11k), pool guide-level counts by perturbation before normalisation, and project onto a gene panel of ≈ 9k genes formed by the union of the top-3,000 HVGs and the on-target genes (Appendix A). We only evaluate models in the perturb-seq + multimodal input settings, replicating the same settings as in the cell-lines described above. Models are evaluated with a 5-fold perturbation-holdout protocol per donor; we report the mean ± std across 2 donors × 5 folds (Table 2). TabPFN is the only model whose top-20 DE predictions are positively correlated with truth on average (cosine +0.108); all other learned models cluster near the Mean baseline (−0.026 cosine).

**Table 2:**
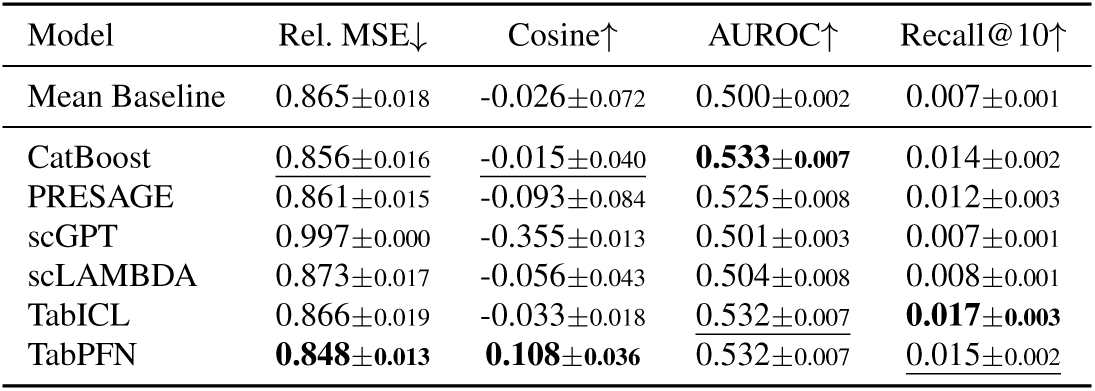
Within-donor prediction on the genome-wide CD4^+^ T cell Perturb-seq screen [Zhu et al., 2025]. Counts are pooled by perturbation before normalization; the gene panel is the union of the top-3,000 HVGs and the perturbations’ on-target genes (≈ 9k genes per donor). Models are evaluated on two donors (D1, D2) with 5-fold perturbation-holdout cross-validation per donor. Values are mean ± std across 2 donors × 5 folds (10 runs per model). Models are trained using Perturb-seq co-expression as well as multimodal representations. Best learned model per column in **bold**, second-best <u>underlined</u>.

| Model | Rel. MSE↓ | Cosine↑ | AUROC↑ | Recall@10↑ |
| --- | --- | --- | --- | --- |
| Mean Baseline | 0.865±0.018 | -0.026±0.072 | 0.500±0.002 | 0.007±0.001 |
| CatBoost | <u>0.856±0.016</u> | <u>-0.015±0.040</u> | **0.533±0.007** | 0.014±0.002 |
| PRESAGE | 0.861±0.015 | -0.093±0.084 | 0.525±0.008 | 0.012±0.003 |
| scGPT | 0.997±0.000 | -0.355±0.013 | 0.501±0.003 | 0.007±0.001 |
| scLAMBDA | 0.873±0.017 | -0.056±0.043 | 0.504±0.008 | 0.008±0.001 |
| TabICL | 0.866±0.019 | -0.033±0.018 | <u>0.532±0.007</u> | **0.017±0.003** |
| TabPFN | **0.848±0.013** | **0.108±0.036** | 0.532±0.007 | <u>0.015±0.002</u> |

Despite TabPFN’s lead, the absolute metrics remain modest compared with the Perturb-seq cell-line benchmarks (Section 4.2). Two factors compound to explain this. (i) *Weak per-perturbation signal from a genome-wide design*: the logFC standard deviation across perturbations is ≈ 0.057 with ∼93% of values within ±0.1 (Appendix Table 3), likely due to higher cell-level variations compared to cell-line screens: primary CD4^+^ T cells are more heterogeneous across donors and batches than immortalised cell lines. (ii) *Pooled guide-level pseudobulk* (∼2 guides per gene) yields substantially noisier targets than cell-line Perturb-seq, which typically pools 30–100+ cells per guide. Overall, these results suggest that for in-vivo perturbation systems stronger emphasis should be put in predictive models that account for cell- and sample-level variability, which is unsurprisingly higher than in immortalized cell-lines.

### 4.3 Zebrafish developmental perturbation atlas

To test whether our findings generalize beyond Perturb-seq transcriptomic readouts, we evaluate on a fundamentally different prediction task: forecasting cell type composition changes during zebrafish embryonic development. The zscape atlas Saunders et al. [2023] profiles ∼2.7M cells via scRNA-seq across 28 genetic perturbations - including single-gene knockouts, combinatorial perturbations, and mutant variants - at 5 developmental timepoints (18, 24, 36, 48, 72 hpf). Cells are annotated into 99 broad cell types; we drop cells in (cell type, timepoint) pairs with ≤ 10 cells (601 cells across 35 pairs, leaving 98 cell types) and formulate the task as: given a perturbation and timepoint, predict the proportion of each cell type across embryos. Per-timepoint dataset statistics (cells, retained cell types, perturbation counts, training-frame rows) are summarised in Appendix Table 4.

This setting differs from the Perturb-seq benchmarks in three key respects: (i) the readout is cell type composition rather than gene expression, (ii) the prediction target is inherently compositional (proportions summing to 1), and (iii) the perturbations act on developmental programs - disrupting cell fate decisions and lineage trajectories - rather than steady-state gene regulation.

#### Cross-validation design

We use a 5-fold *strict timepoint holdout*: each fold removes all rows - perturbed *and* control - at one developmental timepoint (18, 24, 36, 48, 72 hpf in fold order), so the models have never seen any data at the held-out developmental stage. All models receive the same underlying embeddings, derived from control cells only: (i) **intervention embeddings** (300 dimensions): we compute a pseudobulk profile per gene across control cells, select 2,000 HVGs (force-including all perturbation target genes), apply truncated SVD to the genes × cells matrix, and standardize; for combinatorial perturbations, component gene embeddings are averaged; (ii) **cell state embeddings** (300 dimensions): mean expression of 300 HVGs per cell type, standardized across cell types; (iii) **phenotype embeddings** (98 dimensions): control cell type proportions per timepoint, standardized across timepoints. This is in line with the Prophet evaluation Ji et al. [2025].

#### Model configurations

All learned models predict the full 98-dimensional cell-type proportion vector for each (perturbation, timepoint) condition. Prophet Ji et al. [2025] receives the three embedding matrices natively through its intervention - cell state-phenotype decomposition and is trained from scratch with a 2-layer Transformer (see Appendix B).

Tabular regressor configurations (CatBoost / TabICL / TabPFN), the Mean and Bootstrap baselines, and the per-fold cross-validation splits are described in detail in Appendix A.3.

#### Results

The mean baseline reaches *R*^2^ = 0.352 but collapses to *R*^2^ = −0.49 at 18 hpf, the earliest developmental stage and the hardest backward extrapolation (Figure 3). TabPFN is the strongest learned model overall, achieving the best aggregate Spearman correlation (0.808) and *R*^2^ (0.704; Table 5) albeit not significant compared to CatBoost. CatBoost is the next strongest tabular baseline (Spearman 0.763, *R*^2^ = 0.662), Prophet reaches comparable *R*^2^ (0.651) but lower Spearman correlation (0.533), and TabICL remains above the Mean baseline but trails the other learned models on *R*^2^ (0.604) while retaining higher Spearman correlation (0.741) than Prophet. Since all four learned models predict the same 98-dim vector from the same intervention/phenotype embeddings, the gap to the Mean baseline reflects what attention-based or boosted regressors can recover from ∼70 (perturbation, timepoint) training rows; the gap among learned models reflects pretraining priors and architectural choices rather than feature parity. The Pearson curves give the same overall ordering and are reported in Appendix A.3, Figure 4.

**Figure 3:**
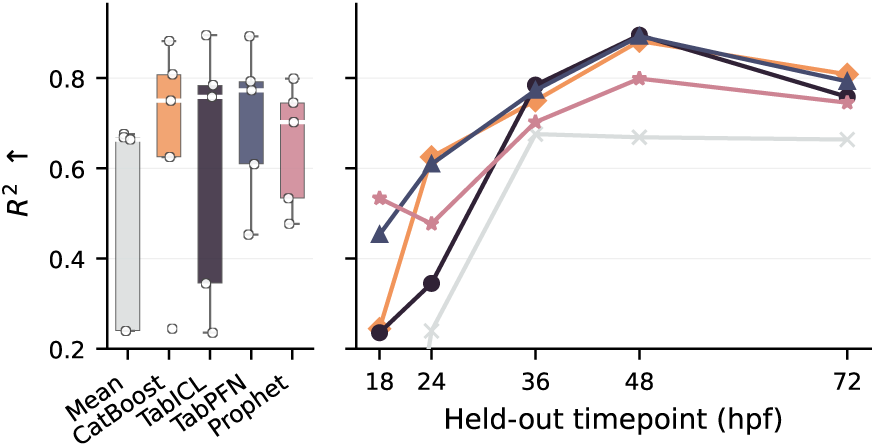
Per-timepoint generalisation under strict time-point holdout, *R*^2^. **Left**: side boxplot of the five fold values per model (colours act as the legend). **Right**: per-fold means; y-axis cut at 0.2 (Mean baseline at 18 hpf, *R*^2^ ≈ −0.5, drops below). Pearson counterpart in Appendix Figure 4.

## 5 Discussion and Conclusion

Across three experimental settings, TabICL and TabPFN match or outperform specialized perturbation prediction models despite using no biology-specific pretraining. A simple recipe of leveraging PCA to decompose the output space, followed by zero-shot tabular regression, is competitive from single-cell cross-cell-type prediction to pseudobulk Perturb-seq and zebrafish developmental composition forecasting. This suggests that, in these benchmarks, strong posterior-predictive regression and effective featurization matter more than architecture-specific biological inductive biases.

Three aspects of the evaluations presented in the manuscript are worth underlying: First, PCA turns high-dimensional response prediction into a collection of well-conditioned scalar regression problems, precisely the regime where prior-fitted tabular models are strongest; our ablations show that the load-bearing property is the *top-variance subspace* of the target rather than the PCA basis itself. Second, the OT matching procedure extends the same componentwise regression idea to the cell level, allowing tabular models to generate realistic target-cell distributions without cell-level pretraining. Third, the zebrafish results further indicate that this formulation transfers beyond gene-expression, enabling to translate predictions to functional phenotypes.

While this study is preliminary — limited to the application and evaluation of PFNs on the cellular perturbation prediction problem — much remains to explore in terms of capabilities such as leveraging and evaluating the entire predictive distribution [Landsgesell et al., 2026] and interpretability [Ye et al., 2025]. A particularly promising direction stems from the fact that PFNs are pre-trained on structural causal models: their in-context representations can be mined to recover the causal structure underlying their predictions [Swelam et al., 2025, Bühler et al., 2026]. Extracting such causal relationships from in-context cellular perturbation data is closely analogous to inferring gene regulatory networks [Kim et al., 2024, Sextro et al., 2026, Neto, 2025], suggesting that PFNs could serve as a basis for causal discovery in biology.

## A Dataset Descriptions

### A.1 Pseudobulk Perturbation Datasets

We evaluate on five pseudobulk datasets derived from two large-scale Perturb-seq studies:

#### Nadig et al. datasets

Two cell lines from Nadig et al.: (i) **nadig_hepg2**—HepG2 hepatocellular carcinoma cells, and (ii) **nadig_jurkat**—Jurkat T-lymphocyte cells. Both contain CRISPR-based perturbations with pseudobulk-aggregated expression profiles stored as .h5ad files.

#### Replogle et al. datasets

Three datasets from Replogle et al.: (i) **replogle_k562_essential**— K562 chronic myelogenous leukemia cells targeting essential genes, (ii) **replogle_k562_gw**—K562 genome-wide perturbation screen, and (iii) **replogle_rpe1_essential**—RPE1 retinal pigment epithelium cells targeting essential genes.

All pseudobulk datasets specify a perturbation column identifying each genetic perturbation, a control label for unperturbed cells, and a batch column encoding experimental covariates.

The splits for the 5 datasets were identical to the one defined by the PRESAGE paper [Littman et al., 2025].

### A.2 CD4 Genome-Wide Dataset

The CD4 genome-wide (GW) dataset is derived from Zhu et al. [2025] and consists of genome-wide CRISPR perturbations in primary human CD4^+^ T cells from two donors (D1=CE0008162, D2=CE0006864). We use the Stim48hr condition only and start from guides passing the keep_for_DE quality filter; non-targeting guides are merged under the label control.

#### Perturbation filter

The provided DE_stats table (Stim48hr) lists 11,281 perturbations. We retain only those with ontarget_significant == True, i.e., perturbations whose knockout significantly downregulated their own target gene; this leaves 7,193 perturbations across the dataset. The remaining ≈ 4k perturbations are excluded from both training and evaluation.

#### Pooled pseudobulk

For each retained donor we (i) restrict the guide-level pseudobulk to that donor’s Stim48hr rows passing keep_for_DE, (ii) pool raw counts by perturbation (one row per perturbation, including a single control row built from all NTC guides), and (iii) apply normalize_total(target_sum=10^4^) ◦ log1p. Pooling before normalisation tightens the per-perturbation target estimate compared with averaging post-normalised guide pseudobulks.

#### Gene panel

Per donor we form the union of (a) the top-3,000 HVGs of the pooled adata and (b) the on-target genes of the 7,193 significant perturbations that appear in the donor’s variable set. This yields ≈ 8.9k genes per donor (D1: 8,918 genes, 6,934 rows; D2: 8,913 genes, 7,002 rows). DEGs are precomputed per perturbation (top 3,000 genes by absolute DESeq2 log-fold-change), restricted to the donor-specific gene panel.

#### Bootstrap oracle

Because each pooled adata has exactly one row per perturbation, the hierarchical resampler used elsewhere as a noise floor (Appendix B.10) collapses to a deterministic oracle (cosine → 1, relMSE → 0). We therefore omit the Bootstrap row from Table 2.

#### Splits

Perturbations are split into five perturbation-disjoint random folds (60%/20%/20% train/val/test) per donor with a fixed seed; control is always assigned to train. After filtering this gives 4,160/1,387/1,386 perturbations per fold for D1 and 4,201/1,400/1,400 for D2. Each model is trained 5 times per donor (2 donors × 5 folds = 10 runs).

#### Modality features

Tabular cell-match models (CatBoost, TabICL, TabPFN) concatenate per-perturbation feature vectors from ten modalities: five Perturb-seq pseudobulk references (Replogle K562 GW, K562 Essential, RPE1 Essential; Nadig HepG2, Jurkat), three protein-text or structure-based priors (ESM2, BioGPT, CellProfiler), two STRING-DB pathway embeddings, and DepMap CRISPR gene-effect scores. A gene co-expression embedding (PCA on the training-fold logFC matrix) is concatenated as an additional perturbation feature by default.

#### Effect-size comparison vs. K562 GW

The modest absolute metrics in Table 2 are dominated by biological signal weakness rather than technical noise. The pooled-pseudobulk pipeline operates on raw guide-level counts (no within-guide normalisation precedes the pooling step), so summing across guides per perturbation is mathematically equivalent to summing all cells of all guides directly, and the median guide carries 49 cells (mean 79, IQR 17–100, max 10,717), giving total cells per perturbation comparable to or higher than typical cell-line Perturb-seq references. Splits are within-donor (see the *Splits* paragraph above), so cross-donor variability does not enter any single evaluation. What does differ is the underlying logFC distribution: Table 3 compares per-perturbation logFC statistics on K562 GW (Replogle) and on CD4 GW D1/D2. K562 GW logFC standard deviation is ≈ 3.2× larger than CD4 GW D2; the fraction of perturbation–gene entries inside ±0.1 jumps from 59.4% on K562 GW to 93.3% on CD4 GW D2. The two donors are within ∼ 10% of each other on every statistic. Because most genome-wide knockouts in primary CD4^+^ T cells produce essentially null transcriptomic responses at the pseudobulk level, very little signal remains for any model to predict.

**Table 3:**
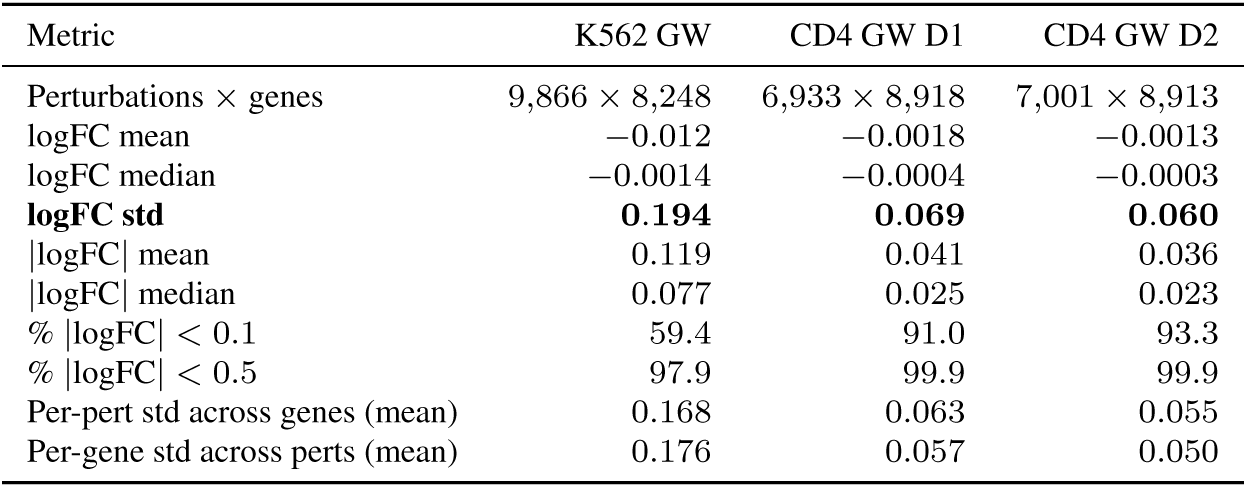
Per-perturbation logFC statistics on K562 GW (Replogle, cached pseudobulk in logFC form) and CD4 GW D1 / D2 (per-perturbation log1p-normalised expression with the control row subtracted as reference). All numbers computed over every perturbation × gene entry of the respective matrix.

### A.3 Zebrafish Developmental Perturbation Atlas (zscape)

The zscape dataset comprises 2,687,135 single cells and 32,031 genes from 812 zebrafish embryos carrying 28 genetic perturbations across 5 developmental timepoints (18, 24, 36, 48, 72 hpf). Perturbations include 18 single-gene knockouts (*cdx4*, *egr2b*, *epha4a*, *foxd3*, *foxi1*, *hand2*, *hgfa*, *hoxb1a*, *mafba*, *met*, *noto*, *phox2a*, *smo*, *tbx1*, *tbx16*, *tbxta*, *tfap2a*, *zc4h2*), 5 combinatorial perturbations (*cdx4-cdx1a*, *tbx16-msgn1*, *tbx16-tbx16l*, *tfap2a-foxd3*, *wnt3a-wnt8*), and 5 mutant variants (*hgfa-mut*, *mafba-mut*, *met-mut*, *noto-mut*, *tbx16-mut*). Six control conditions (*ctrl-inj*, *ctrl-hgfa*, *ctrl-mafba*, *ctrl-met*, *ctrl-noto*, *ctrl-tbx16*) are unified under a single control label, contributing 610,804 cells (23% of the dataset). Cells are annotated into 99 broad cell types; we drop cells in (cell type, timepoint) pairs with ≤ 10 cells (601 cells across 35 pairs), removing one cell type entirely (*mineralized tissue, bone*) and leaving 98 cell types in the final vocabulary.

#### Data processing

The raw scRNA-seq counts are processed as follows: (i) control labels are unified; (ii) rare (cell type, timepoint) pairs are dropped as described above, retaining 2,686,534 cells; (iii) for each of the 812 embryos, we count the number of cells belonging to each of the 98 retained cell types and normalize by total cell count to obtain per-embryo cell type proportions; (iv) proportions are averaged across embryo replicates within each (perturbation × timepoint) condition, yielding 86 conditions (81 perturbed + 5 control timepoints). Per-timepoint statistics—number of profiled cells, retained cell types, measured perturbations, and rows in the training DataFrame—are summarised in Table 4; the totals are skewed across stages (252K cells / 15 perturbations at 18 hpf vs. 813K cells / 24 perturbations at 36 hpf), which directly maps onto per-fold difficulty under strict timepoint holdout (Figure 3). The full training DataFrame contains 7,775 rows, where each row corresponds to one (perturbation, timepoint, cell type) triplet—restricted to retained (cell type, timepoint) pairs—with the cell type proportion as the target value (ranging from 0 to ∼0.04). In parallel, an embryo-level pseudobulk expression matrix (812 × 32,031) is constructed by summing counts per embryo, normalizing to 10^4^ counts, and applying log1p—this is used only for embedding construction, not as a prediction target.

**Table 4:** Per-timepoint statistics for the zscape dataset after rare-pair filtering. Cells profiled per timepoint and the number of retained cell types reflect the underlying biology (more cell types appear at later developmental stages), while perturbation coverage is uneven (most perturbations were measured at 18–48 hpf, only six at 72 hpf). Under 5-fold strict timepoint holdout, fold *f* holds out timepoint *T_f_* and the entire “Rows” column for that timepoint moves to validation.

| Timepoint | Cells (profiled) | Cell types (retained) | Perturbations | Rows (= cell types × conditions) |
| --- | --- | --- | --- | --- |
| 18 hpf | 252,482 | 87 | 15 | 1,392 |
| 24 hpf | 369,283 | 87 | 19 | 1,740 |
| 36 hpf | 813,341 | 92 | 24 | 2,300 |
| 48 hpf | 786,652 | 94 | 17 | 1,692 |
| 72 hpf | 464,926 | 93 | 6 | 651 |
| Total | 2,686,534 | 98 (union) | 28 (union) | 7,775 |

#### Cross-validation splits

We use a 5-fold *strict timepoint holdout*: each fold *f* ∈ {0*, . . .,* 4} removes *all* rows—perturbed and control alike—at one developmental timepoint, training only on rows at the remaining four timepoints. Fold *f* holds out timepoint *T_f_* ∈ {18, 24, 36, 48, 72} hpf. Per-fold sizes (train rows / val rows / val perturbations) are:

- Fold 0 (18 hpf): 6,383 / 1,392 / 15
- Fold 1 (24 hpf): 6,035 / 1,740 / 19
- Fold 2 (36 hpf): 5,475 / 2,300 / 24
- Fold 3 (48 hpf): 6,083 / 1,692 / 17
- Fold 4 (72 hpf): 7,124 / 651 / 6

Held-out perturbation counts per fold reflect the unequal coverage across timepoints: most perturbations were measured at 36 hpf (24 perturbations), only 6 at 72 hpf. The validation set contains perturbations the model *has* seen at other timepoints, but never at the held-out one; controls at the held-out timepoint are also in val, so the model has zero training rows at the held-out developmental stage. The protocol is therefore much stricter than perturbation-family holdout (which keeps both controls and other perturbations at the held-out timepoint in train, allowing the per-(cell type, time-point) mean to be well estimated and pushing trivial baselines to Spearman ≈ 0.95); strict timepoint holdout breaks this leak entirely and forces the model to extrapolate compositional dynamics to a developmental stage absent from training.

#### Embedding construction

All embeddings are derived from control cells only, ensuring no leakage of perturbation-specific information:

- **Intervention embeddings** (30 × 300): We normalize and log-transform control cells, select 2,000 HVGs (force-including all perturbation target genes), transpose the expression matrix to genes × cells, and apply truncated SVD (*d* = 300, explaining the top variance components). Each perturbation target gene is represented by its SVD coordinates; for combinatorial perturbations, component gene vectors are averaged; for mutant variants, the base gene vector is used. Two zero-vector entries (negative_gene, negative_drug) serve as the control intervention embedding.
- **Cell state embeddings** (98 × 300): Mean expression of 300 HVGs per cell type across control cells, standardized (zero mean, unit variance) across cell types.
- **Phenotype embeddings** (5 × 98): Control cell type proportions per timepoint, standardized across timepoints.

#### Tabular model features and prediction regime

Each tabular regressor (CatBoost, TabICL, TabPFN) processes one row per (perturbation, timepoint) condition—70–80 training rows per fold under strict holdout. Features concatenate the intervention (300 dim) and phenotype (98 dim) embeddings (398-dim total); the cell-state embedding is *not* used as an input feature because all 98 cell types are predicted jointly rather than queried one at a time. The target is the 98-dimensional cell-type proportion vector for that (perturbation, timepoint), with cells absent from a given time-point (the rare-pair-filtered combinations) filled with 0 to maintain a consistent output dimension across rows. Both X and Y are independently PCA-projected to *d_x_* = *d_y_* = 32 components, retaining *>*99% of variance in both spaces given the small training set. CatBoost is fit as a single multi-output regressor on the PCA-transformed Y with the MultiRMSE loss (iterations=1000, depth=6, l2_leaf_reg=3.0); TabICL (n_estimators=8, GPU) and TabPFN (default settings with ignore_pretraining_limits=True, GPU) are fit per Y component (32 univariate fits per fold, since neither natively supports multi-output regression). Predictions in PCA-Y space are inverse-PCA-transformed back to the 98-dim proportion space and scattered to the long format for evaluation.

#### Mean baseline

The Mean baseline predicts, for each validation row, the mean training proportion at the same (cell type, timepoint) pair; under strict timepoint holdout no training rows exist at the validation timepoint, so the prediction falls back to the per-cell-type marginal mean over the four training timepoints (a final fallback to the global mean handles unseen cell types).

#### Evaluation metrics

We evaluate using rank correlation, Pearson correlation, and *R*^2^ score. For each (perturbation × timepoint) condition, metrics are computed over the cell type proportion values present at that timepoint after rare-pair filtering (between 87 and 94 cell types, depending on timepoint; predicted vs. ground truth). The aggregate zscape table reports Spearman correlation and *R*^2^ averaged over the five strict timepoint-holdout folds. Per-fold *R*^2^ is plotted in Figure 3 (main text), and the Pearson counterpart is plotted in Figure 4; the in-text averages are unweighted means over the 5 folds.

**Figure 4:**
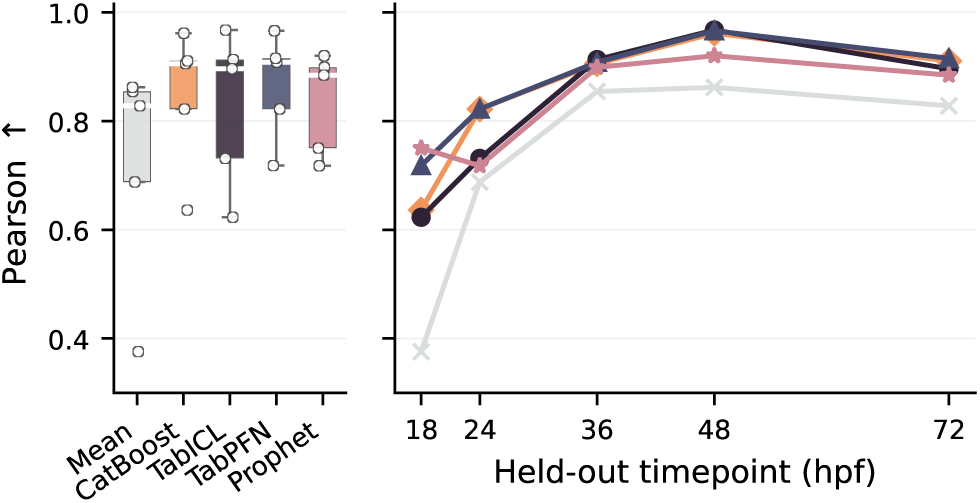
Per-timepoint Pearson under strict timepoint holdout (Pearson counterpart of Figure 3). **Left**: side boxplot of the five fold values per model. **Right**: per-fold mean Pearson across (perturbation, timepoint) conditions. The Pearson curves give the same ordering as *R*^2^ in the main text—TabPFN strongest overall, learned models converging at 36–72 hpf—but with a less dramatic baseline collapse at 18 hpf (Pearson 0.38 vs. *R*^2^ ≈ −0.49).

### A.4 OpenProblems Cell-Level Dataset

The cell-level benchmark uses data from the NeurIPS 2023 OpenProblems chemical-perturbation challenge. The raw dataset (sc_counts.h5ad) contains 298,087 cells from three donors (Donor_1, Donor_2, Donor_3) across 21,265 genes (full transcriptome), annotated into four cell types (T cells, NK cells, B cells, Myeloid cells) and exposed to 144 drug conditions (143 active compounds plus DMSO vehicle control).

**Table 5:** Zebrafish cell type proportion prediction (zscape) under *strict timepoint holdout*: 5-fold CV in which all rows (perturbed and control) at one developmental timepoint are removed per fold (folds 0–4 hold out 18, 24, 36, 48, 72 hpf respectively), so the model has never seen any data at the held-out stage. Each model predicts the full 98-dimensional cell-type proportion vector for one (perturbation, timepoint) condition; tabular regressors use a 398-d feature vector (intervention 300-d + phenotype 98-d) and PCA-decomposed targets at *d_x_* = *d_y_* = 32. Values are mean ± std across the 5 folds. Best per column in **bold**.

| Model | Spearman↑ | $R^2\uparrow$ |
| --- | --- | --- |
| Mean Baseline | 0.596±0.109 | 0.352±0.452 |
| Bootstrap | 0.595±0.109 | 0.347±0.465 |
| CatBoost | 0.763±0.067 | 0.662±0.225 |
| TabICL | 0.741±0.091 | 0.604±0.262 |
| TabPFN | **0.808±0.072** | **0.704±0.155** |
| Prophet | 0.533±0.138 | 0.651±0.124 |

#### Gene panel

Following the STACK paper evaluation protocol, we restrict to 2,000 highly variable genes selected via scanpy.pp.highly_variable_genes(n_top_genes=2000, flavor=seurat_v3) on the raw counts of the full cohort. For each donor we materialise three HDF5 files:

- {Donor}_hvg2k.h5ad — *log-normalised* counts (log_1_ *p* of per-cell counts scaled to 10^4^), 2k HVGs, all cell types and drugs. Consumed by the cell-match tabular models.
- {Donor}_hvg2k_context.h5ad — *raw* counts, T cells only (all drugs), 2k HVGs. The STACK “context.”
- {Donor}_hvg2k_test.h5ad — *raw* counts, DMSO-treated NK/B/Myeloid cells only, 2k HVGs. The STACK “test” / target control pool.

#### Evaluation protocol

All models solve the same cross-cell-type prediction task: given drug-treated source (T cell) profiles and DMSO-treated target cells, predict the distribution of drug-treated target cells under each of the 143 active drugs, separately for NK, B, and Myeloid cell types. Models are run independently per donor (no cross-donor pooling) and evaluated against the ground-truth drug-treated target cells in the same donor.

### A.5 Cell-Level Cross-Cell-Type Pipeline

#### Cell-level prediction via cell matching

A central methodological contribution of this work is a procedure for turning a pseudobulk-style tabular regressor into a single-cell perturbation predictor. Starting from log-normalised expression of source (T cell) and target (NK/B/Myeloid) populations on 2k HVGs, we (i) fit a shared PCA on the combined source and target control cells, (ii) pair source with target control cells in PCA space via optimal transport (OT) with uniform marginals, (iii) train a per-component regressor—TabICL, TabPFN, or CatBoost—on the matched control pairs, and (iv) apply the regressor to drug-perturbed source cells and invert the PCA to obtain cell-level log-normalised target predictions. We benchmark this pipeline against the STACK foundation model [Dong et al., 2026], which consumes raw counts from a source context and a target test set and generates raw-count predictions zero-shot; we align both families on the same 2k-HVG evaluation panel and evaluate with the cell-eval suite.

This section gives end-to-end input/output, normalisation, and evaluation details for the four cell-level models (TabICL, CatBoost, TabPFN with OT cell matching, and STACK).

#### Tabular cell-match models (TabICL / TabPFN / CatBoost)

All three share the same pipeline and differ only in the per-PCA-component regressor:

1. **Input.** A single log-normalised donor file {Donor}_hvg2k.h5ad. We extract three cell populations: *source control* (*S*^ctrl^, T cells with DMSO), *source drug* (*S^d^*, T cells with each of the 143 active drugs), and *target control* (*T* ^ctrl^, DMSO cells of each target type *c* ∈ {NK, B, Myeloid}).
2. 2. **Shared PCA.** For each target cell type *c*, fit sklearn.decomposition.PCA with *k* = 50 components on the row-stacked matrix [*S*^ctrl^; *T _c_*^ctrl^]; project all three populations plus *S^d^* into the same *k*-dim subspace.
3. **OT cell matching.** Compute the |*S*^ctrl^| × |*T_c_*^ctrl^| squared-Euclidean cost matrix in PCA space (normalised by its max for numerical stability), solve exact OT via POT’s ot.emd under uniform marginals, and extract the nonzero entries as source-target index pairs. If |*S*^ctrl^| exceeds 2,000, we subsample source cells uniformly (seed 42) before forming the cost matrix.
4. **Per-component regression.** Let *X* ∈ ℝ*^N^*^×^*^k^* be matched source-control PCA scores and *Y* ∈ ℝ*^N^*^×^*^k^* matched target-control PCA scores. Standardise *X* with a StandardScaler. Then:

- **TabICL** (tabicl.TabICLRegressor, tabicl-regressor-v2-20260212): fits one regressor per output component (*j* = 1*, . . ., k*) with KV-cache enabled for efficient reuse across components; 16 estimators, robust normalisation, outlier threshold 3.5.
- **TabPFN** (tabpfn.TabPFNRegressor): identical per-component loop with 8 estimators, softmax temperature 0.9.
- **CatBoost** (catboost.CatBoostRegressor, loss=MultiRMSE): a single model predicting all *k* components jointly; 1,000 iterations, depth 6, *ℓ*_2_ leaf reg 3.0.
5. **Prediction & inverse PCA.** Apply the fitted regressor to PCA-projected source-drug cells to obtain target PCA scores, then invert the PCA transform to get log-normalised expression over 2k HVGs.
6. **Output.** One HDF5 per drug, with rows concatenated across the three target cell types; obs carries predicted_cell_type and sm_name; X stores log-normalised predicted expression (clipped at 0) in a sparse matrix over 2k genes.

Two variants are reported: **OT matching** (the default, as above) and a **nearest-neighbour (NN) matching** ablation that replaces step 3 with sklearn’s 1-NN between source and target control cells in the same PCA space. All other steps are identical.

#### STACK foundation model

We run the STACK bc_large_aligned checkpoint via the stack-generation command-line binary [Dong et al., 2026] with:

- **Context** {Donor}_hvg2k_context.h5ad: raw counts from source (T cell) cells across all drugs.
- **Test** {Donor}_hvg2k_test.h5ad: raw counts from DMSO-treated target cells (NK, B, Myeloid).
- **Gene list** basecount_1000per_15000max.pkl: STACK’s fixed 15,012-gene vocabulary. The intersection with the 2k HVG panel is what ultimately enters the evaluation.
- **Split column** sm_name: instructs STACK to generate one output per drug.

STACK performs zero-shot in-context generation: for each drug *d* it uses the source-drug and source-control cells as context and the target DMSO cells as the conditioning target, producing raw-count predictions for the target population under drug *d*. The output is written as one HDF5 per drug, with raw counts across the STACK vocabulary; rows inherit cell-type annotations from the test file. At evaluation time we log-normalise STACK outputs (normalize_total(1e4)+log1p) and intersect their variable names with the 2k HVG panel so that all models are compared on an identical gene set.

#### Cell-eval protocol

Evaluation is performed with the cell-eval library’s MetricsEvaluator running with the full profile (CPU, 16 threads). For each (donor, target cell type) pair we pair the predicted AnnData with the real drug-treated target cells from the same donor, restrict both to the intersection of variable names, filter to the intersection of drugs actually predicted, and compute all metrics per drug. The headline table (Table 1) reports mean ± SEM across the 3 target cell types (NK, B, Myeloid) for Donor 1, the donor for which all four models—including TabPFN with OT matching—have completed runs; the OT-vs-NN comparison in Table 10 aggregates across all 3 donors × 3 target cell types for TabICL and CatBoost. The subsampled context-cell-count sweep (Figure 5) uses the same pipeline with reduced source populations at fractions {0.1, 0.25, 0.5, 0.75, 1.0} and PCA *k* = 50.

## B Model Details

### B.1 TabICLv2

We use the tabicl-regressor-v2-20260212 checkpoint with n_estimators = 8 and x_pca_n_components =y_pca_n_components = 128. The model is applied zero-shot without fine-tuning and processes inputs in batches of 128.

### B.2 TabPFN

TabPFN [Hollmann et al., 2025] is the original Prior-Fitted Network for tabular regression. We use TabPFN 2.6 (tabpfn.TabPFNRegressor) with n_estimators = 8 and ignore_pretraining_limits=True; the same PCA decomposition pipeline as TabICLv2 is used (*d* = 128 components for output spaces and *d* = 64 for input spaces, with one regressor fit per principal component). Like TabICLv2, the model is applied zero-shot without fine-tuning.

### B.3 CatBoost

We train CatBoost gradient-boosted decision trees with iterations = 1000, depth = 6, and l2_leaf_reg = 3.0. On CatBoost Multi-output regressor is trained for each fold split and experiment, using the same PCA decomposition pipeline as TabICLv2.

### B.4 PRESAGE

PRESAGE is trained with gene representations are constructed from *n* = 128 NMF embedding dimensions. The model uses node2vec-derived network embeddings, GAT-based pathway weighting, and over 40 embedding modalities including GenePT, BioGPT, ESM, CellProfiler morphological features, StringDB interaction scores, DepMap dependency profiles, and MSigDB pathway memberships. For the pseudobulk cell-lines runs, we used the best configurations as shared in the PRESAGE original repository, so models were trained with the best set of hyperparameters as defined by the authors, for each dataset and split. For the CD4 T Cells benchmark, we reused the hyperparameters of the K562 GW experiment, given that the number of genes and perturbations was similar.

### B.5 scGPT

scGPT uses the whole_human pretrained checkpoint. We fine-tune with batch_size = 64, max_epochs = 20, and learning rate lr = 1 × 10^−4^.

### B.6 scLAMBDA

scLAMBDA is trained for training_epochs = 200 with batch_size = 500, learning rate lr = 5 × 10^−4^, latent_dim = 30, and hidden_dim = 512.

### B.7 STACK

STACK uses the bc_large_aligned pretrained checkpoint and performs zero-shot inference without fine-tuning. It operates on the BaseCount gene vocabulary and is evaluated on the cell-eval benchmark.

### B.8 Prophet

Prophet is a Transformer-based model that decomposes perturbation experiments into three components—intervention, cell state, and phenotype—each processed through learned tokenizer networks before being fed as a sequence to a Transformer encoder. The model predicts a scalar readout (here, a cell type proportion) from the encoder’s output.

#### Architecture

We train Prophet from scratch on the zscape dataset (no pretrained checkpoint). The architecture (646K parameters) processes inputs as follows:

1. **Tokenization.** Each embedding is projected to the model dimension via a 2-layer MLP: Linear(*d*_in_) → GELU → Dropout(0.2) → Linear(128). This yields separate tokenizer networks for interventions (*d*_in_ = 300), cell state (*d*_in_ = 300), and phenotype (*d*_in_ = 98).
2. **Sequence assembly.** With simpler = true, the Transformer input is a 3-token sequence: a learnable CLS token plus two intervention tokens (for iv1 and iv2). Cell state and phenotype embeddings are injected into the regression head rather than the Transformer sequence.
3. **Transformer encoder.** 2-layer TransformerEncoder with model_dim = 128, 1 attention head, feedforward dimension 256, and dropout 0.1.
4. **Regression head.** The CLS token output is passed through a 2-layer MLP: 128 → GELU → Dropout(0.2) → 128 → GELU → 1.

#### Training

Loss: MSE. Optimizer: AdamW (lr = 10^−4^, weight decay 0.01). Schedule: cosine warmup (5,000 warmup steps, 50,000 max steps). Batch size: 512. Dropout: 0.2 for all tokenizer and regressor layers. Early stopping: patience 10 on validation *R*^2^ (hardcoded in Prophet). Training converges in ∼66–74 epochs (∼25 minutes per fold on a single NVIDIA A40 GPU).

### B.9 Mean Baseline

The Mean baseline predicts the mean training treatment effect (perturbation mean minus control mean) for every unseen perturbation, serving as a negative control.

### B.10 Bootstrap Baseline and Experimental Noise Floor

#### Implementation

The Bootstrap baseline estimates the experimental noise floor via hierarchical bootstrapping. For each perturbation in the validation/test set, the procedure is:

1. Let *B* be the set of batches containing both perturbed and control cells for that perturbation.
2. **Outer resampling**: sample |*B*| batches from *B* with replacement.
3. **Inner resampling**: within each resampled batch, sample cells with replacement from the perturbed population and (independently) from the control population, matching the original cell counts.
4. Compute pseudobulk profiles by averaging expression across the resampled cells (separately for perturbed and control), then compute the treatment effect 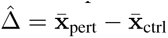.

This produces a single noisy replicate of the true treatment effect per perturbation. Running with different random seeds via –multirun yields the full bootstrap distribution. Our datasets are log-normalized (max expression values 5–7), so we use mean aggregation rather than sum-normalize-log1p pseudobulk computation.

#### Relationship to prior work

Our bootstrap is conceptually equivalent to the experimental error estimate introduced by Cole et al. [2026], who perform the same hierarchical resampling but computes *M* bootstrap replicates and reports the 90th percentile of the error distribution 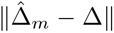 as the experimental error bound *ϵ*(*q*=0.9). Our implementation differs in framing: we evaluate a single bootstrap replicate as a “prediction” using the same evaluation suite as all other models, rather than computing a quantile threshold. The underlying measurement—how much the treatment effect estimate varies due to finite cell sampling—is identical.

#### Why the bootstrap serves as an oracle upper bound

Since the Bootstrap has access to the held-out perturbation’s own cells, it achieves near-perfect performance on all metrics. On relative MSE restricted to the top-20 differentially expressed genes, the Bootstrap achieves values of 0.007–0.017 across datasets (Table 6), while the best learned model (TabICL or TabPFN) achieves 0.38–0.73. On cosine similarity (top-20 DE), the Bootstrap reaches 0.97–0.99 while learned models reach 0.22–0.58.

**Table 6:** Bootstrap vs. best learned model on relative MSE (top-20 DE). The Bootstrap uses the held-out perturbation’s own cells (oracle access), while learned models must generalize from perturbation embeddings alone.

| Dataset | Bootstrap | Best model | Gap (×) |
| --- | --- | --- | --- |
| HepG2 | 0.008 | 0.468 | 59× |
| Jurkat | 0.010 | 0.483 | 48× |
| K562 Essential | 0.011 | 0.506 | 46× |
| K562 Genome-Wide | 0.017 | 0.729 | 43× |
| RPE1 Essential | 0.007 | 0.377 | 54× |

## C Evaluation Metrics

### C.1 Pseudobulk Metrics

All the pseudobuk metrics are derived from PRESAGE [Littman et al., 2025].

#### Relative MSE (top-*k* DE)

For each perturbation *p*, let G*_k_*^(*p*)^ denote the set of *k* genes with the largest absolute differential expression. The relative MSE is:

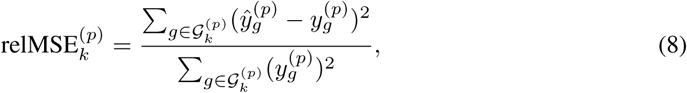

where *y_g_*^(*p*)^ is the ground-truth control-subtracted expression and 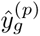 is the prediction. Values below 1.0 indicate the model outperforms the zero-vector (mean control) baseline.

#### Cosine similarity (top-*k* DE)

Cosine similarity between predicted and ground-truth expression vectors restricted to the top-*k* DE genes:

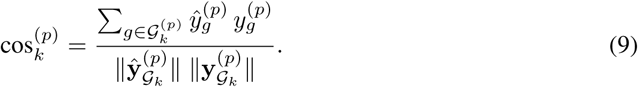

#### Phenocopy AUROC

For each training perturbation *i*, ground-truth neighbors are defined as perturbations whose cosine similarity exceeds the threshold *θ* = median(*S*) + *m* · MAD(*S*), where *S* is the vector of all pairwise cosine similarities and MAD is the median absolute deviation. The AUROC is computed using predicted cosine similarities as scores against these binary neighbor labels.

#### Phenocopy Recall@*k*

The fraction of ground-truth top-*k* neighbors (by cosine similarity in the true expression space) that also appear among the top-*k* neighbors in the predicted space.

### C.2 Cell-Level Metrics

All the pseudobuk metrics are derived from STATE [Adduri et al., 2025].

#### Pearson delta

Pearson correlation between the mean expression change (treated minus control) per drug in the predicted versus ground-truth profiles.

#### DE Spearman

Spearman rank correlation computed on statistically significant differentially expressed genes.

#### DE direction match

Fraction of significant DE genes for which the predicted direction of change (up- or downregulation) matches the ground truth.

#### DE significant gene recall

Recall of the set of significant DE genes: the fraction of true significant genes that are also predicted as significant.

#### Discrimination score

Cosine-similarity-based separability of treated cells from control cells, measuring whether predicted perturbation responses are distinguishable from background variation.

#### Pearson E-distance

Implemented as pearson_edistance in the cell-eval library. For each perturbation *p*, we compute the (squared-Euclidean) energy distance 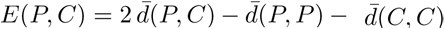 between perturbed cells *P* and control cells *C* (where 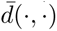 is the mean pairwise distance), separately on the real and the predicted data, yielding two per-perturbation vectors **e**_real_, **e**_pred_ ∈ ℝ*^N^*^pert^ . The reported metric is the Pearson correlation *ρ*(**e**_real_, **e**_pred_) ∈ [−1, 1], so higher is better; this is a correlation between two vectors of distances rather than a distance itself.

## D Experimental Setup

### D.1 Cross-Validation Protocol

All pseudobulk and CD4 experiments use 5-fold cross-validation with perturbation-aware splits (random seed 42). Control perturbations are always assigned to the training set. Non-control perturbations are divided across folds such that each perturbation appears in exactly one test fold.

### D.2 Data Efficiency Sweep

To assess sample efficiency, we evaluate models at training fractions {0.25, 0.5, 0.75, 1.0} of the available training perturbations. This sweep is applied to all three pseudobulk models (TabICLv2, PRESAGE, CatBoost) across all five pseudobulk datasets, yielding 3 × 5 × 4 × 5 = 300 experimental configurations (models × datasets × fractions × folds).

### D.3 Cell-Level Context Sweep

For the cell-level benchmark, we vary the fraction of source (T cell) training data at {0.1, 0.25, 0.5, 0.75, 1.0}. This sweep is applied to TabICLv2 (with optimal transport alignment), CatBoost (with optimal transport alignment), and STACK, all evaluated on 5,000 HVG features.

### D.4 Perturb-seq + Multi-Modal vs. Multi-Modal Only

We compare two feature regimes for the pseudobulk perturbation-prediction pipeline. In both regimes the model receives a single concatenated feature vector per perturbation; the regimes differ only in whether Perturb-seq pseudobulk references are included.

- **Perturb-seq + multi-modal** (the default; “Perturb-seq” in figure / table labels): for each perturbation *g* we look up its pseudobulk profile in every Perturb-seq dataset *other than* the held-out target dataset (e.g., when the target is Nadig HepG2 we use Replogle K562 Essential, K562 GW, RPE1 Essential, and Nadig Jurkat as references); each reference profile is reduced to a fixed-rank PCA representation and the four per-reference vectors are concatenated. We then concatenate the multi-modal embeddings (ESM2 protein embeddings, BioGPT text embeddings, CellProfiler morphological features, two STRING-DB pathway-interaction embeddings, and DepMap CRISPR gene-effect scores) onto the same vector. Perturbations missing in a given reference contribute a zero block. The resulting feature vector is the model input; the prediction target is the held-out dataset’s pseudobulk profile.
- **Multi-modal only** (“multi-modal” in figure / table labels): the same concatenated multi-modal embeddings as above, but without the Perturb-seq references. This isolates the contribution of same-modality (gene-expression) features versus auxiliary modalities.

The Perturb-seq + multi-modal regime is the default reported throughout the paper and in Figure 2 and Table 8; the multi-modal-only ablation appears in Figure 7 and Table 9.

### D.5 Compute Resources

All pseudobulk and CD4 experiments were run on a single NVIDIA L40 or A40 GPU. STACK inference required 128 GB RAM, 8 CPUs, and up to 6 hours of walltime per donor.

#### Per-fold wallclock for the pseudobulk benchmark

Table 7 reports the median per-fold wallclock for the four models we logged end-to-end (training + inference) on the five Perturb-seq datasets. Because TabICL and TabPFN are pretrained Prior-Fitted Networks applied *zero-shot*, their per-fold wallclock is just the inference pass over the held-out perturbations (∼3 s on a single GPU). CatBoost trains and predicts within 4–7 s per fold on its boosted-tree multi-output regressor. PRESAGE, in contrast, is trained from scratch each fold (10,000 epochs by default) and takes approximately two orders of magnitude longer (median ≈ 253 s, max ≈ 272 s on the Replogle Essential datasets). The overall picture is that *pretraining moves the cost outside the per-experiment loop*: TabICL and TabPFN match the simplest gradient-boosted baseline on raw walltime while producing accuracy comparable to or better than the much more expensive trained-from-scratch PRESAGE.

**Table 7:** Per-fold wallclock on the pseudobulk perturbation benchmark, aggregated across 5-fold CV and the five Perturb-seq datasets (Nadig HepG2/Jurkat, Replogle K562 Essential / K562 GW / RPE1 Essential). Values are median seconds per fold, measured on a single NVIDIA L40 or A40 GPU. “Pretraining” denotes how the model arrives at inference time: zero-shot PFNs amortise their cost into pretraining, while PRESAGE is trained from scratch per fold.

| Model | Pretraining | Per-fold wallclock (s, median) |
| --- | --- | --- |
| TabICL | zero-shot PFN | 3 |
| TabPFN | zero-shot PFN | 3 |
| CatBoost | trained per fold | 4 |
| PRESAGE | trained per fold | **253** |

## E Additional Figures

**Figure 5:**
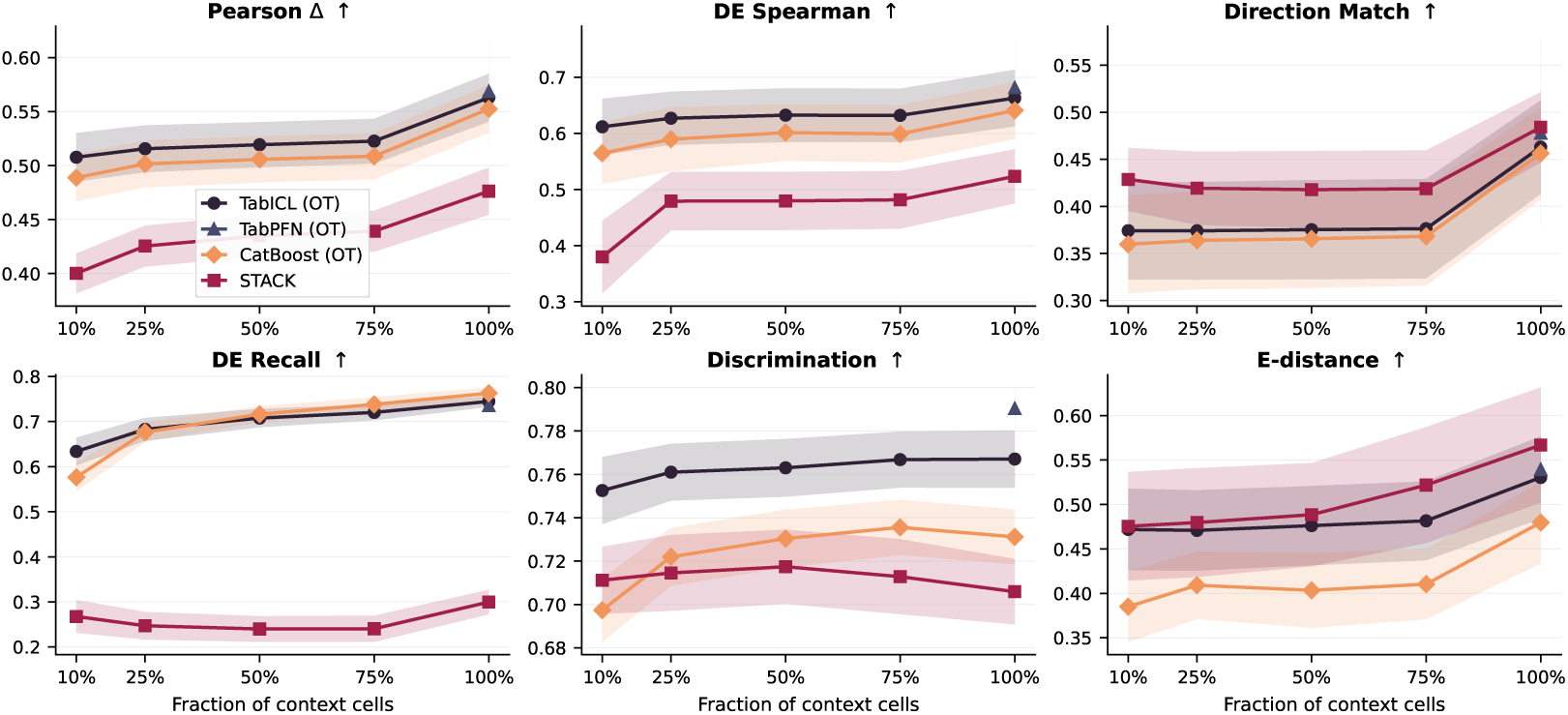
Context cell count sweep for cell-level prediction. TabICL (OT) and CatBoost (OT) degrade gracefully, while STACK shows greater sensitivity to context size.

## F Ablation Studies

## G Mechanism Analyses: Why Do TFMs Work?

Tabular foundation models (TFMs) such as TabICL and TabPFN consistently match or outperform specialised perturbation-prediction baselines in our benchmarks (Section 4.2), yet it is not *a priori* clear which component of the pipeline produces this advantage. We test two natural candidate mechanisms: (i) in-context inference that exploits local structure of the training support set, and (ii) a PCA-based feature geometry that happens to make the perturbation regression approximately linear. These correspond, respectively, to the support-set ablation in §G.1 (mechanism i) and the output-geometry ablation in §G.2 (mechanism ii).

**Figure 6:**
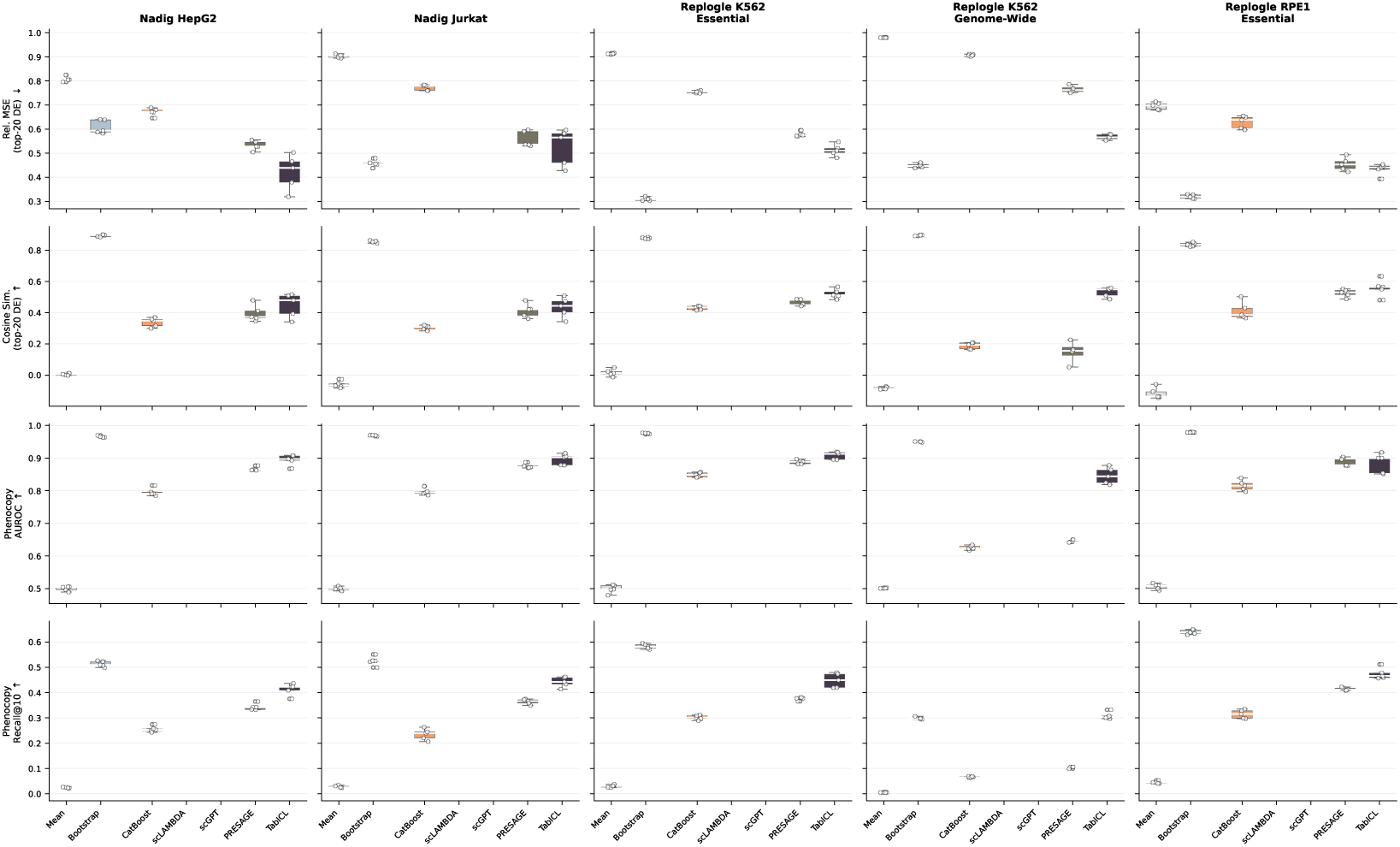
Pseudobulk perturbation prediction in the default *Perturb-seq + multi-modal* feature regime, across five Perturb-seq datasets and four evaluation metrics. Each boxplot summarizes 5-fold CV results for one model on one dataset. Mean (negative control) and Bootstrap (experimental noise estimate) are separated from learned models by a gap. The Bootstrap serves as an oracle upper bound with access to the held-out perturbation’s cells (see Appendix B.10). Among learned models, TabICL and TabPFN consistently achieve the best performance. Models concatenate Perturb-seq pseudobulk references from non-held-out cell lines with multi-modal embeddings (Appendix D.4); the multi-modal-only ablation (Figure 7) drops the Perturb-seq references and degrades performance.

Each study isolates one mechanism by holding the other fixed: the support-set ablation varies the training context but keeps the feature pipeline and the model architecture fixed; the geometry ablation varies the output *y*-decomposition with controlled interventions (tail-PCA, random orthogonal rotation, random Gaussian projection, ICA, NMF) while keeping the context and the per-component regressor fixed.

### G.1 Diversity-aware support-set selection

#### Motivation and hypothesis

In-context regressors operate by conditioning on an in-context support set at inference time. Two candidate mechanisms can explain their strong perturbation-prediction performance: the *size* of the support set (more examples, stronger prior) or its *composition* (the specific perturbations and how well they cover the response manifold). If composition matters, then actively selecting a diverse subset of training perturbations as the support set should recover most of the full-context performance from a small fraction of the pool — and in some regimes may even exceed full context, because redundant near-duplicate perturbations are downweighted. We test this directly.

#### Data and splits

Three Perturb-seq datasets (Nadig HepG2, Nadig Jurkat, Replogle RPE1 Essential) and both TFMs (TabICL, TabPFN) with 5-fold perturbation-disjoint splits (seed 42) shared with the headline pseudobulk benchmark.

**Table 8:** Pseudobulk perturbation prediction in the default *Perturb-seq + multi-modal* feature regime: per-perturbation Perturb-seq pseudobulk references from non-held-out cell lines are concatenated with multi-modal embeddings (ESM2, BioGPT, CellProfiler, STRING-DB, DepMap; see Appendix D.4) and fed to the regressor. Values are mean ± std across 5-fold CV. Best per row in **bold**.

| Dataset | Mean Baseline | Bootstrap | CatBoost | PRESAGE | scGPT | scLAMBDA | TabPFN | TabICL |
| --- | --- | --- | --- | --- | --- | --- | --- | --- |
| *Rel. MSE*↓ |  |  |  |  |  |  |  |  |
| Nadig HepG2 | 0.807±0.011 | **0.008±0.000** | 0.612±0.011 | 0.530±0.010 | 0.969±0.008 | 0.678±0.008 | 0.472±0.016 | 0.468±0.015 |
| Nadig Jurkat | 0.903±0.007 | **0.010±0.001** | 0.714±0.010 | 0.586±0.025 | 0.944±0.005 | 0.755±0.017 | 0.491±0.033 | 0.483±0.032 |
| Replogle K562 Essential | 0.913±0.002 | **0.011±0.001** | 0.697±0.003 | 0.569±0.009 | 0.948±0.007 | 0.738±0.013 | 0.506±0.009 | 0.508±0.011 |
| Replogle K562 GW | 0.980±0.001 | **0.017±0.001** | 0.856±0.005 | — | 0.990±0.001 | 0.919±0.013 | 0.729±0.010 | 0.731±0.010 |
| Replogle RPE1 Essential | 0.696±0.015 | **0.007±0.000** | 0.578±0.023 | 0.456±0.017 | 0.970±0.003 | 0.635±0.037 | 0.382±0.020 | 0.377±0.021 |
| *Cosine*↑ |  |  |  |  |  |  |  |  |
| Nadig HepG2 | 0.005±0.007 | **0.995±0.001** | 0.423±0.024 | 0.387±0.034 | -0.078±0.028 | 0.191±0.025 | 0.456±0.027 | 0.458±0.023 |
| Nadig Jurkat | -0.061±0.022 | **0.992±0.000** | 0.401±0.030 | 0.398±0.051 | 0.042±0.030 | 0.193±0.021 | 0.451±0.035 | 0.459±0.031 |
| Replogle K562 Essential | 0.015±0.023 | **0.990±0.000** | 0.485±0.007 | 0.468±0.017 | 0.030±0.025 | 0.239±0.014 | 0.502±0.012 | 0.510±0.007 |
| Replogle K562 GW | -0.084±0.008 | **0.973±0.001** | 0.222±0.019 | — | 0.059±0.013 | 0.220±0.011 | 0.240±0.014 | 0.222±0.017 |
| Replogle RPE1 Essential | -0.117±0.036 | **0.990±0.001** | 0.494±0.031 | 0.558±0.031 | -0.505±0.030 | 0.141±0.056 | 0.570±0.024 | 0.583±0.017 |
| *AUROC*↑ |  |  |  |  |  |  |  |  |
| Nadig HepG2 | 0.498±0.008 | **0.984±0.002** | 0.876±0.004 | 0.863±0.008 | 0.496±0.005 | 0.698±0.010 | 0.880±0.006 | 0.879±0.006 |
| Nadig Jurkat | 0.499±0.004 | **0.987±0.001** | 0.892±0.003 | 0.874±0.005 | 0.498±0.008 | 0.706±0.012 | 0.905±0.004 | 0.903±0.006 |
| Replogle K562 Essential | 0.500±0.006 | **0.989±0.001** | 0.900±0.009 | 0.888±0.008 | 0.492±0.012 | 0.749±0.010 | 0.908±0.008 | 0.908±0.007 |
| Replogle K562 GW | 0.502±0.003 | **0.975±0.001** | 0.644±0.006 | — | 0.503±0.005 | 0.568±0.003 | 0.647±0.008 | 0.647±0.008 |
| Replogle RPE1 Essential | 0.501±0.010 | **0.993±0.000** | 0.894±0.008 | 0.878±0.017 | 0.508±0.014 | 0.698±0.023 | 0.910±0.009 | 0.904±0.011 |
| *Recall@10*↑ |  |  |  |  |  |  |  |  |
| Nadig HepG2 | 0.024±0.002 | **0.619±0.013** | 0.331±0.009 | 0.339±0.016 | 0.022±0.005 | 0.167±0.014 | 0.349±0.012 | 0.350±0.011 |
| Nadig Jurkat | 0.026±0.004 | **0.651±0.017** | 0.343±0.016 | 0.356±0.022 | 0.024±0.004 | 0.170±0.011 | 0.376±0.018 | 0.375±0.016 |
| Replogle K562 Essential | 0.030±0.004 | **0.666±0.006** | 0.367±0.010 | 0.384±0.010 | 0.027±0.006 | 0.213±0.013 | 0.389±0.013 | 0.391±0.011 |
| Replogle K562 GW | 0.005±0.001 | **0.402±0.005** | 0.088±0.003 | — | 0.007±0.001 | 0.044±0.003 | 0.105±0.003 | 0.106±0.002 |
| Replogle RPE1 Essential | 0.041±0.001 | **0.775±0.009** | 0.420±0.008 | 0.397±0.014 | 0.034±0.005 | 0.204±0.014 | 0.461±0.009 | 0.460±0.010 |

#### Support-set selection strategies

For every (model, dataset, fold) we subsample the training perturbations to pct_train ∈ {0.25, 0.5, 0.75} using two strategies:

- **Random**: uniform sample with a fixed seed (42). The same seed is used across folds so that fold-to-fold variance reflects data variance, not sampling variance.
- **FPS (farthest-point sampling on pseudobulk effect vectors)**: for each training perturbation *p* we form the effect vector 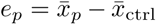 (pseudobulk log-fold-change), reduce to 16 PCs via PCA fit on the training effect matrix, seed with the perturbation of largest ∥*e_p_*∥, then iteratively expand *S* by arg max*_p_* min*_q_*_∈_*_S_ ∥e_p_* − *e_q_*∥_2_ until |*S*| = ⌈pct_train · |train|⌉. Fully deterministic given (fold, pct_train).

FPS explicitly maximises coverage of the response manifold; random is an unbiased reference. The 100% baseline is the full training set, reused from the pseudobulk headline sweep without rerun. We run 5 folds per configuration, giving 6 (model,dataset) × 2 strategies × 3 pcts × 5 folds = 180 runs, plus 30 shared baseline runs.

#### Results

FPS helps both TFMs on RPE1 Essential and Jurkat, with the gain over random largest at the smallest context fraction we tested (pct_train = 0.25); on HepG2 the two strategies are essentially indistinguishable across all fractions (Table 11, Figure 12). On RPE1 Essential at pct_train = 0.25, FPS raises cosine(top-20 DE) by +0.127 for TabICL (0.503 → 0.630) and +0.195 for TabPFN (0.416 → 0.611), *exceeding* the 100% baseline (0.583 / 0.570) with only a quarter of the training perturbations. The effect is intermediate on Jurkat (+0.04–0.05 cosine at 25%) and absent or slightly negative on HepG2 (−0.02 to 0 cosine). Relative MSE shows the same pattern: FPS lowers relMSE by up to −0.08 (TabPFN RPE1 at 25%). At pct_train = 0.75 the two strategies converge on HepG2 and Jurkat; the RPE1 gap narrows but FPS is still ahead (+0.04–0.05 cosine).

On the datasets where FPS helps (RPE1 Essential, Jurkat), the TFMs benefit substantially from **diversity-aware curation** of the in-context training set: selecting a maximally spread subset of perturbation effect vectors—via a cheap deterministic FPS pass—recovers most of the full-context signal from a small fraction of the training pool, and on RPE1 surpasses the full-context baseline because redundant, near-duplicate perturbations drop out of the context. The effect is strongest when (i) the context is small relative to the response manifold and (ii) the training pool contains many low-effect perturbations that a diversity-aware selector correctly downweights; HepG2 satisfies neither condition strongly, which is consistent with the absence of an FPS gain there.

**Figure 7:**
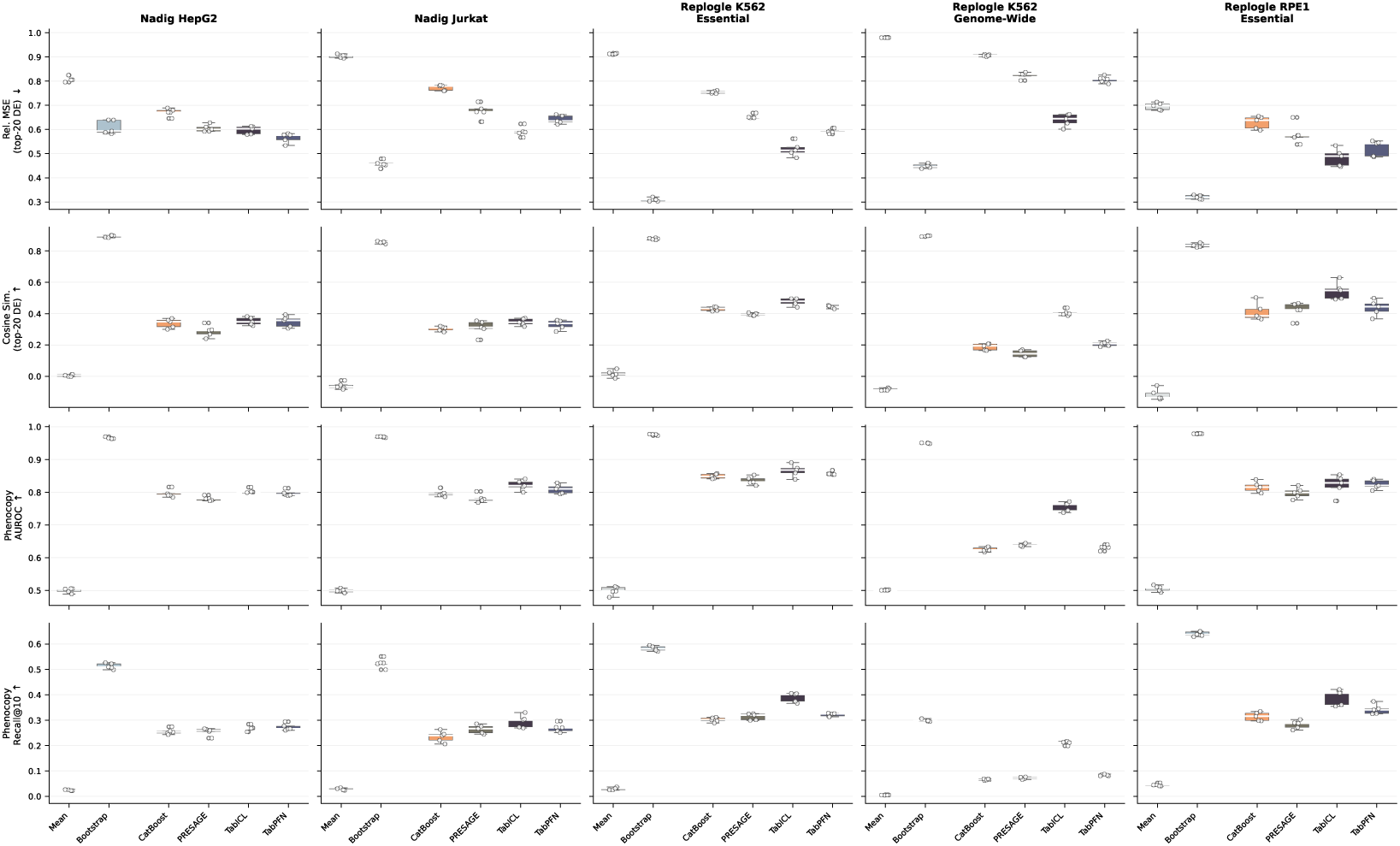
Pseudobulk perturbation prediction in the *multi-modal only* feature regime, across five Perturb-seq datasets and four evaluation metrics. Same layout as Figure 6 but the Perturb-seq pseudobulk references are dropped from the input; models see only the multi-modal embeddings (ESM2, BioGPT, CellProfiler, STRING-DB, DepMap; see Appendix D.4). TabPFN and TabICL are included alongside CatBoost and PRESAGE; scGPT and scLAMBDA are excluded as they operate on raw Perturb-seq expression and do not consume multi-modal features. Removing the Perturb-seq references degrades performance relative to the default regime in Figure 6, indicating that same-modality (gene-expression) features carry the bulk of the signal.

### G.2 Output-space geometry ablation

#### Motivation and hypothesis

The TabICL/TabPFN multiview pipeline projects the gene-space target onto a *d_y_* = 128 PCA basis and fits one regressor per component. Replacing this PCA basis with alternatives that differ in a single controlled way lets us attribute the perturbation-prediction performance to specific geometric properties of the output space rather than to “PCA in general”. We test the hypothesis:

> **H_geom_.** TFM performance depends on (i) *which* 128-dimensional subspace the gene-space target is projected into and (ii) the non-orthogonality of that basis, but is approximately invariant to (iii) an orthogonal rotation of the basis within a fixed subspace.

The three subclaims pin down exactly what property of PCA is load-bearing. (i) is tested by pca_tail (bottom-128 PCs — same rank, opposite end of the variance spectrum) and gaussian_projection (random 128-d Gaussian projection — same rank, unrelated subspace). (ii) is tested by nmf (a non-orthogonal, non-negative basis in a related subspace). (iii) is tested by pca_random_rot (PCA followed by a fixed random orthogonal rotation, same subspace and same Parseval identity) and ica (FastICA on the top-128 PCA subspace, a principled rotation that maximises per-axis non-Gaussianity).

**Table 9:** Pseudobulk perturbation prediction in the *multi-modal only* feature regime: the Perturb-seq pseudobulk references are dropped and the regressor sees only the multi-modal embeddings (ESM2, BioGPT, CellProfiler, STRING-DB, DepMap; see Appendix D.4). Values are mean ± std across 5-fold CV. Best per row in **bold**.

| Dataset | Mean Baseline | Bootstrap | CatBoost | PRESAGE | TabPFN | TabICL |
| --- | --- | --- | --- | --- | --- | --- |
| *Rel. MSE*↓ |  |  |  |  |  |  |
| Nadig HepG2 | 0.807±0.011 | **0.008±0.000** | 0.673±0.016 | 0.613±0.015 | 0.566±0.020 | 0.549±0.018 |
| Nadig Jurkat | 0.903±0.007 | **0.010±0.001** | 0.773±0.012 | 0.681±0.027 | 0.641±0.017 | 0.608±0.022 |
| Replogle K562 Essential | 0.913±0.002 | **0.011±0.001** | 0.754±0.006 | 0.652±0.011 | 0.592±0.009 | 0.571±0.009 |
| Replogle K562 GW | 0.980±0.001 | **0.017±0.001** | 0.908±0.003 | – | 0.806±0.014 | 0.805±0.016 |
| Replogle RPE1 Essential | 0.696±0.015 | **0.007±0.000** | 0.631±0.025 | 0.572±0.047 | 0.523±0.032 | 0.494±0.040 |
| *Cosine*↑ |  |  |  |  |  |  |
| Nadig HepG2 | 0.005±0.007 | **0.995±0.001** | 0.333±0.031 | 0.311±0.014 | 0.348±0.036 | 0.345±0.022 |
| Nadig Jurkat | -0.061±0.022 | **0.992±0.000** | 0.305±0.014 | 0.299±0.052 | 0.330±0.030 | 0.332±0.028 |
| Replogle K562 Essential | 0.015±0.023 | **0.990±0.000** | 0.431±0.013 | 0.406±0.017 | 0.443±0.008 | 0.442±0.013 |
| Replogle K562 GW | -0.084±0.008 | **0.973±0.001** | 0.190±0.022 | – | 0.206±0.015 | 0.197±0.015 |
| Replogle RPE1 Essential | -0.117±0.036 | **0.990±0.001** | 0.397±0.047 | 0.435±0.058 | 0.437±0.050 | 0.467±0.044 |
| *AUROC*↑ |  |  |  |  |  |  |
| Nadig HepG2 | 0.498±0.008 | **0.984±0.002** | 0.795±0.012 | 0.777±0.002 | 0.797±0.009 | 0.795±0.011 |
| Nadig Jurkat | 0.499±0.004 | **0.987±0.001** | 0.796±0.009 | 0.780±0.013 | 0.810±0.014 | 0.804±0.015 |
| Replogle K562 Essential | 0.500±0.006 | **0.989±0.001** | 0.847±0.008 | 0.837±0.009 | 0.858±0.005 | 0.855±0.007 |
| Replogle K562 GW | 0.502±0.003 | **0.975±0.001** | 0.625±0.005 | – | 0.631±0.007 | 0.632±0.005 |
| Replogle RPE1 Essential | 0.501±0.010 | **0.993±0.000** | 0.813±0.014 | 0.788±0.030 | 0.824±0.014 | 0.824±0.019 |
| *Recall@10*↑ |  |  |  |  |  |  |
| Nadig HepG2 | 0.024±0.002 | **0.619±0.013** | 0.253±0.010 | 0.250±0.007 | 0.273±0.013 | 0.276±0.014 |
| Nadig Jurkat | 0.026±0.004 | **0.651±0.017** | 0.235±0.023 | 0.261±0.018 | 0.269±0.017 | 0.270±0.021 |
| Replogle K562 Essential | 0.030±0.004 | **0.666±0.006** | 0.297±0.011 | 0.304±0.018 | 0.322±0.007 | 0.327±0.008 |
| Replogle K562 GW | 0.005±0.001 | **0.402±0.005** | 0.065±0.004 | – | 0.083±0.003 | 0.082±0.003 |
| Replogle RPE1 Essential | 0.041±0.001 | **0.775±0.009** | 0.312±0.017 | 0.283±0.031 | 0.342±0.020 | 0.347±0.025 |

**Table 10:** Optimal-transport (OT) vs nearest-neighbour (NN) cell matching on the OpenProblems 2k HVG benchmark. Mean across 3 donors × 3 target cell types. OT matching outperforms NN for both regressors (TabICL, CatBoost). TabPFN is omitted because the OT runs only completed for Donor 1.

| Model | Pearson $\Delta$↑ | DE Spearman↑ | Direction↑ | DE Recall↑ | Discrim.↑ | Pearson E-dist.↑ |
| --- | --- | --- | --- | --- | --- | --- |
| TabICL (OT) | **0.563** | **0.663** | **0.463** | **0.744** | **0.767** | **0.530** |
| TabICL (NN) | 0.490 | 0.602 | 0.426 | 0.696 | 0.713 | 0.435 |
| CatBoost (OT) | **0.552** | **0.641** | **0.456** | **0.763** | **0.731** | **0.480** |
| CatBoost (NN) | 0.441 | 0.520 | 0.401 | 0.641 | 0.664 | 0.454 |

#### Decomposition interventions

The six conditions are summarised in Figure 9 (baseline pca plus five alternatives). All five alternatives keep the rank fixed at *d_y_* = 128, keep the input side of the pipeline (*x*-decomposition, context, regressor hyperparameters, PRESAGE-style evaluation) identical to the pseudobulk headline sweep, and intervene only on the *y*-decomposition step.

#### Data, splits, and runs

Same three Perturb-seq datasets (Nadig HepG2, Nadig Jurkat, Replogle RPE1 Essential) and same 5-fold perturbation-disjoint splits as §G.1. Two TFMs (TabICL, TabPFN). Six configs × 5 decomposition alternatives × 5 folds = 150 new runs; the pca baseline (30 runs) is reused from the pseudobulk headline sweep.

#### Results

Figure 9 reports cosine(top-20 DE) mean ± SEM per (model, dataset, decomposition); each panel’s dashed line marks the PCA baseline. The six-row mean Δ relative to the PCA baseline decomposes as follows (in order of magnitude):

**Figure 8:**
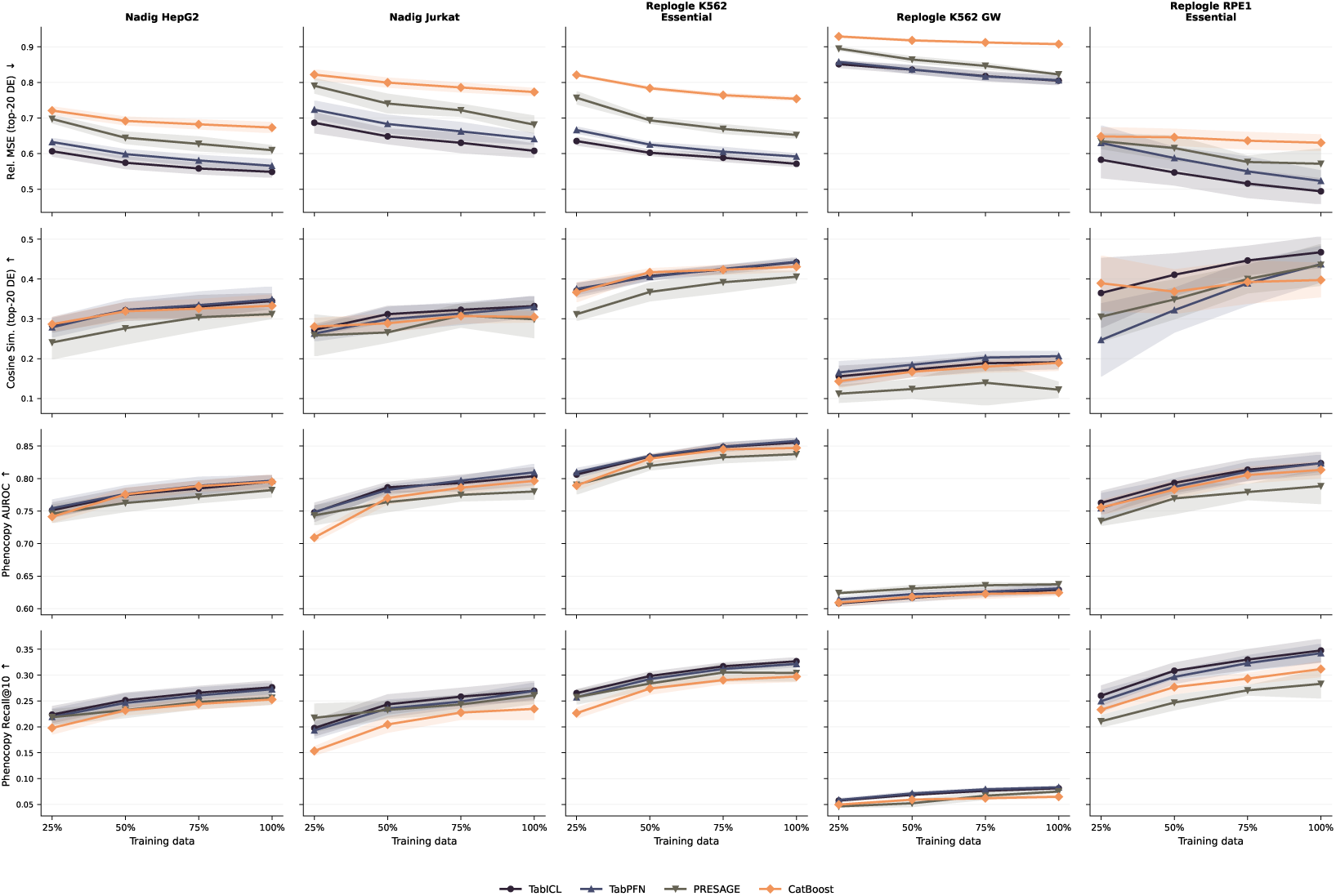
Data efficiency across five Perturb-seq datasets. Each panel shows model performance as a function of training data fraction (25–100%). The Tabular Foundation Models (TabICL, TabPFN) maintain competitive performance even with limited training data. Shaded regions indicate ±1 standard deviation across folds.

- **Subspace matters (large negative** Δ). pca_tail collapses all six rows to near-zero cosine 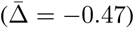: the tail of the variance spectrum carries essentially no perturbation-prediction signal. gaussian_projection retains some signal (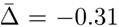, end values 0.12–0.20) but loses most of it: a random 128-d subspace is far weaker than the variance-maximising top-128. Together these pin down *which 128-d subspace* as a dominant factor.
- **Rotation within the subspace is approximately a no-op (near-zero Δ).** pca_random_rot averages 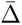 = +0.019, consistent with the theoretical Parseval-based null. ica averages 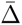 = −0.024 (a small but visible drop), with TabPFN more sensitive to it than TabICL; this is the residual non-linearity in *y* leaking through.
- **Non-orthogonal NMF is, surprisingly, positive (**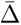 = +0.117). The non-orthogonal NMF basis *improves* cosine on every (model, dataset) row, with the largest gains on TabPFN Jurkat (+0.19) and TabPFN HepG2 (+0.19). NMF with Parseval-breaking non-negative constraints is competitive and in several cases preferable to PCA on the current pipeline. We flag this as a follow-up direction rather than a main claim because (a) relmse_top20_de does not track cosine on NMF as cleanly, hinting at a scale or sign issue, and (b) NMF’s non-negative shift interacts with the per-component robust normaliser in a way that we have not fully diagnosed.

#### Interpretation

The data support H_geom_ on subclaims (i) and (iii) and partially contradict (ii): the TFM’s advantage is robustly tied to the *top-variance subspace of the target* (projecting onto the tail or a random subspace is catastrophic), but is essentially invariant to any orthogonal rotation within that subspace (PCA ≈ random rotation ≈ ICA to within 0.05 cosine). Non-orthogonal NMF does not hurt and in fact helps — contrary to what we expected, and worth further investigation. Combined with the support-set result in §G.1, the geometric picture is that **PCA is not a uniquely required decomposition; what matters is that the 128-d target subspace aligns with the directions of largest variance in the training effect matrix.**

**Figure 9:**
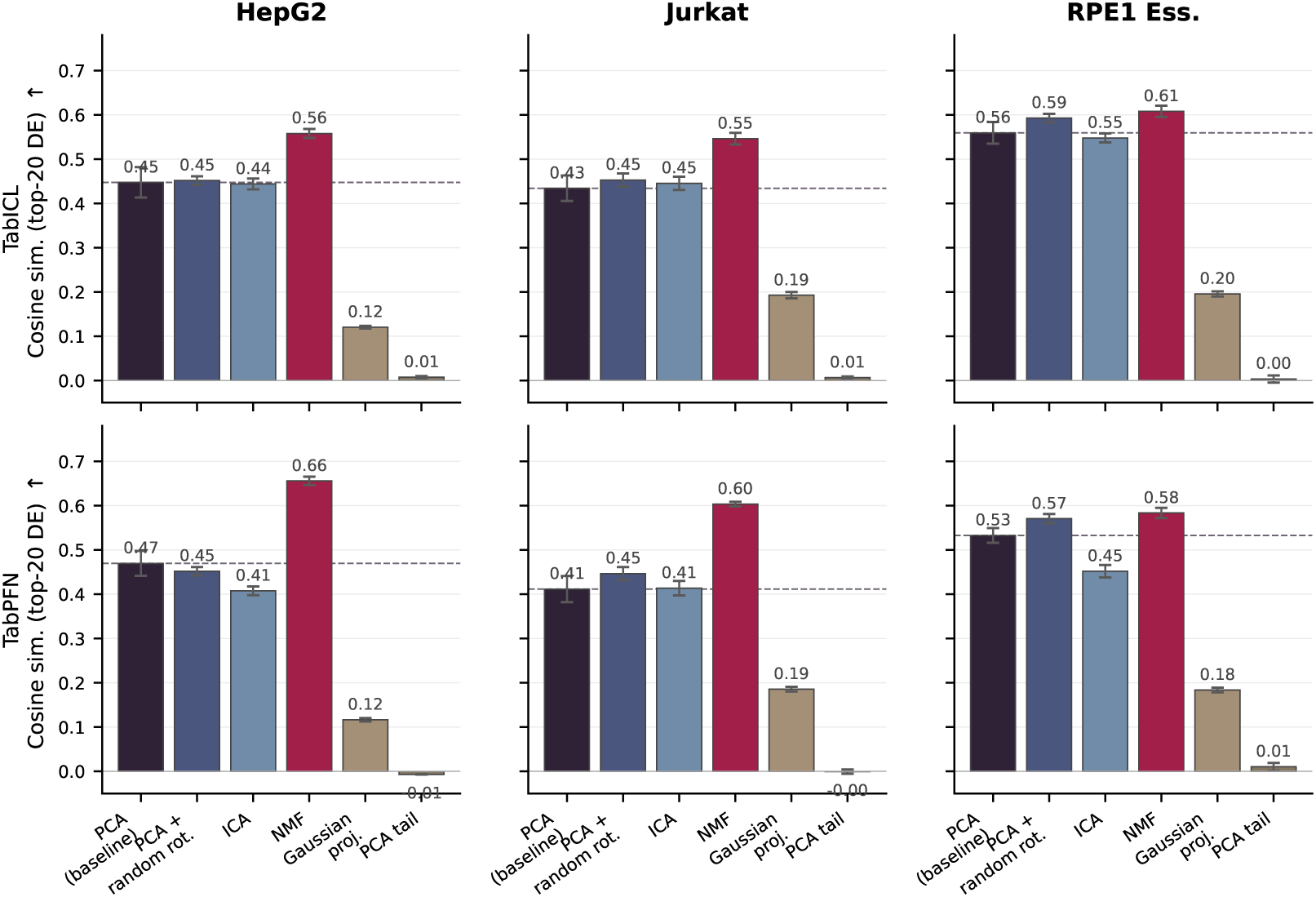
Output-space decomposition ablation. Each panel shows cosine similarity (top-20 DE) for the six *y*-decomposition variants of the TabICL/TabPFN multiview pipeline, with the input pipeline, context, and per-component regression held fixed. PCA (baseline) is reused from the pseudobulk headline sweep; the dashed line in each panel marks the PCA baseline value. PCA + random rot. applies a random orthogonal rotation to the top-128 PCA basis (preserves subspace and Parseval identity); ICA is FastICA on the same top-128 PCA subspace; NMF uses 128 non-negative components (non-orthogonal); Gaussian proj. replaces the subspace with a random 128-d Gaussian projection; PCA tail keeps the bottom-128 PCs. Bars are mean ± SEM across 5 folds. Subspace-breaking interventions (Gaussian proj., PCA tail) collapse cosine to near zero; within-subspace rotations (PCA + random rot., ICA) leave it essentially unchanged; NMF is a positive deviation. Detailed methodology and per-cell numbers in Appendix G.2.

**Figure 10:**
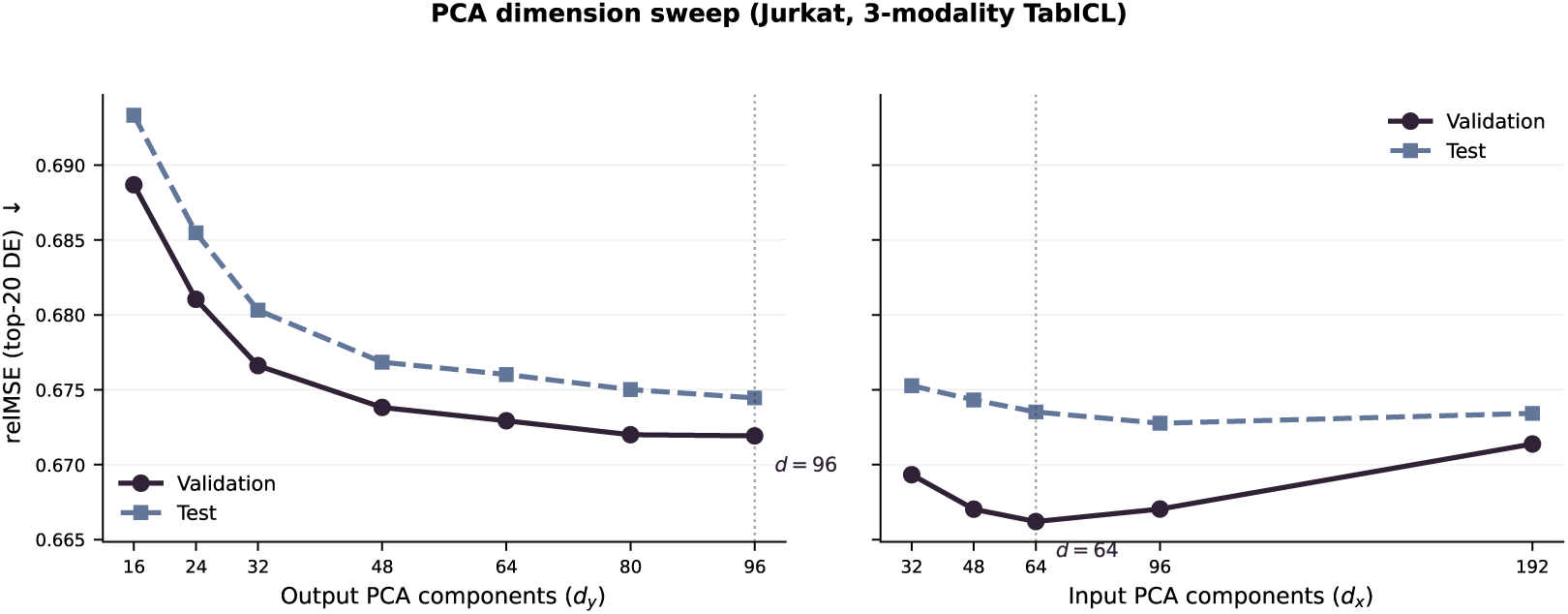
Effect of PCA dimensionality on TabICL performance (Jurkat, 3-modality model). **Left:** Output PCA components (*d_y_*), controlling the dimensionality of the gene expression target space. Performance improves monotonically up to *d_y_* ≈ 80–96, with diminishing returns beyond *d_y_* = 48. **Right:** Input PCA components (*d_x_*), controlling the dimensionality of the perturbation feature space. Optimal at *d_x_* = 64, with degradation at higher dimensions due to noise amplification.

**Figure 11:**
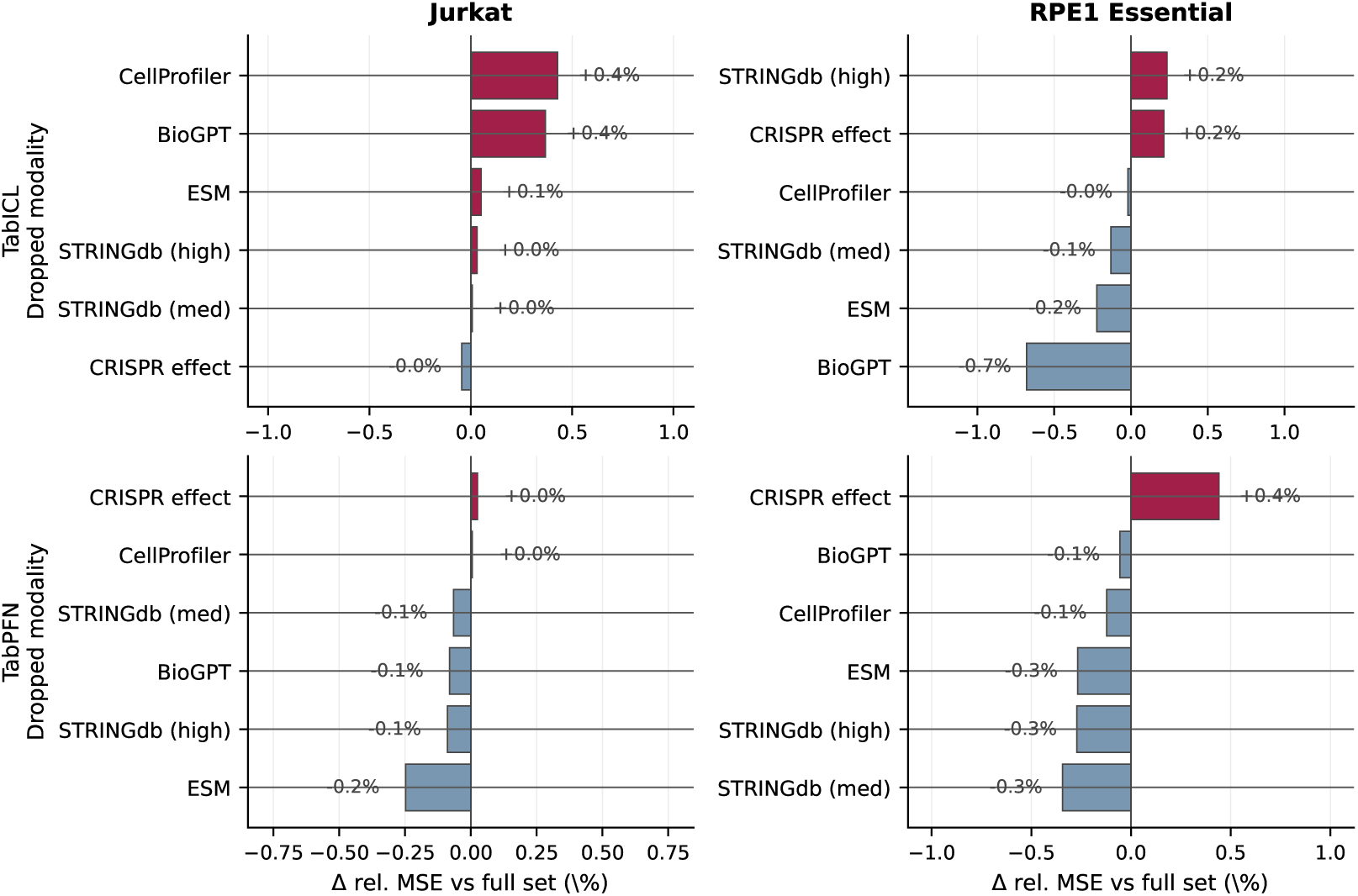
Leave-one-out modality ablation. Each panel shows the percentage change in relative MSE (top-20 DE) when one of the six top embedding modalities (ESM, BioGPT, CellProfiler, STRINGdb-high, STRINGdb-medium, DepMap CRISPR) is dropped from the TabICL / TabPFN multiview model, relative to the full-modality baseline. Bars are mean ± SEM across 5 perturbation-disjoint folds; positive values indicate the modality is helpful (dropping it increases error). Models are run in the perturbseq variant (top modalities concatenated with cross-cell-line Perturb-seq pseudobulk features), so each modality is one of many features and individual drops produce small effects (|ΔrelMSE| *<* 1% in most cells). The most impactful drops differ across (model, dataset) pairs and are not uniformly dominated by any single modality family.

We also probe whether the TFMs are sensitive to the specific choice of in-context training perturbations. Replacing random subsampling with farthest-point sampling (FPS) on pseudobulk effect vectors - a selection strategy designed to maximise coverage of the response manifold - improves performance at reduced context sizes, with the effect most pronounced on the smallest context fraction we tested (Appendix G.1, Figure 12 and Table 11). On RPE1 Essential at pct_train = 0.25, FPS lifts cosine top-20 DE by +0.127 (TabICL) and +0.195 (TabPFN) over random, and remarkably matches or exceeds the 100% baseline with only a quarter of the training perturbations. This indicates that TFMs benefit from a diversity-aware support set: curating a maximally spread subset of training perturbations allows the models to recover most of the full-context signal from substantially fewer in-context examples.

**Figure 12:**
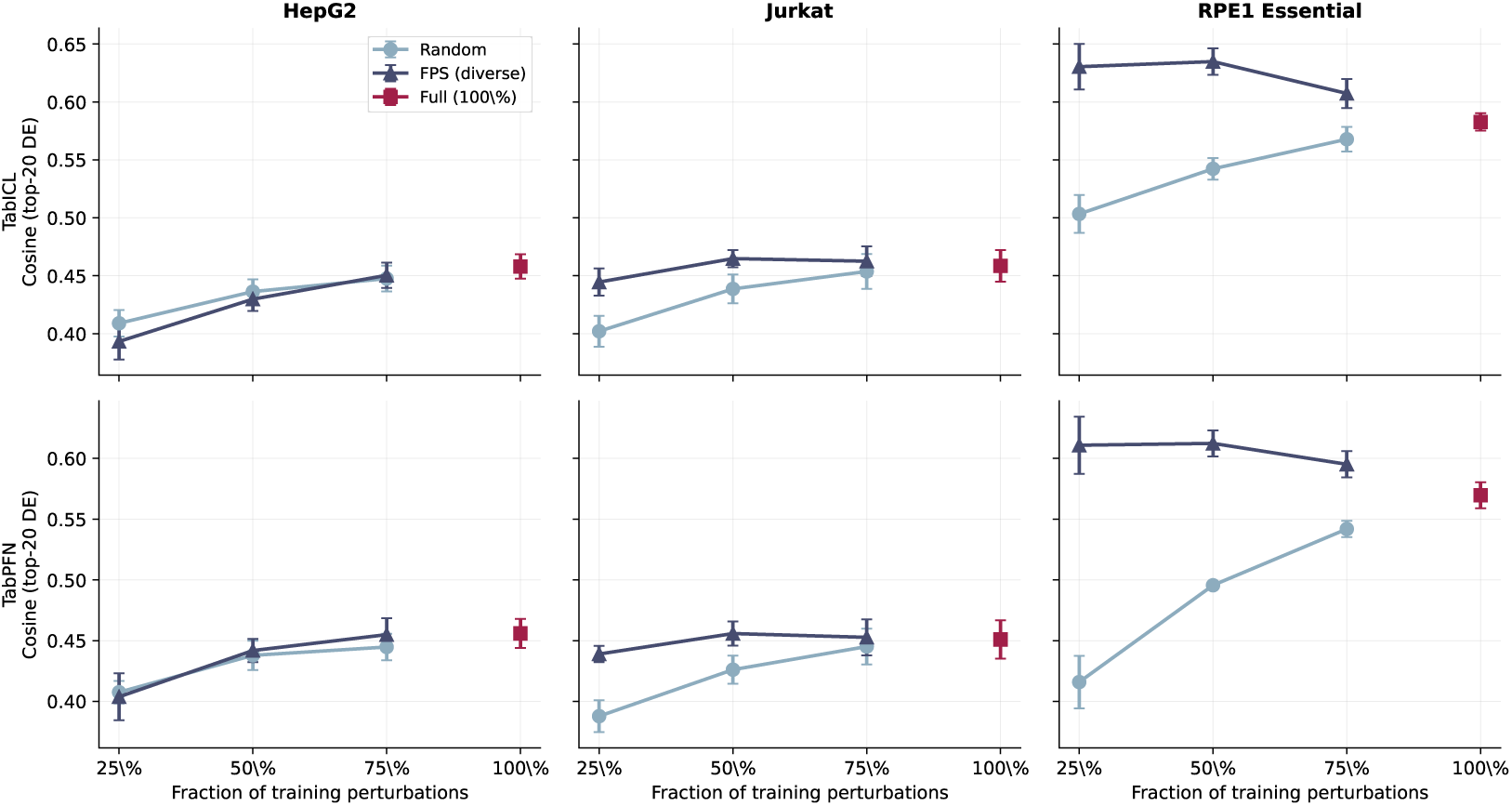
Support-set ablation: cosine similarity (top-20 DE) vs fraction of training perturbations on 3 Perturb-seq datasets × 2 TFMs. Random (grey circles) and FPS (blue triangles) overlap at every fraction; the red square shows the 100% baseline. Error bars are SEM across 5 folds.

**Table 11:** Support-set ablation for TabICL and TabPFN. Each row shows cosine similarity (top-20 DE) at a given fraction of training perturbations, comparing *random* subsampling (same seed across folds) with *farthest-point sampling (FPS)* on pseudobulk effect vectors. *Full* is the 100% baseline. Mean ± SEM across 5 folds.

| Dataset | Frac. | Random | FPS | Full |
| --- | --- | --- | --- | --- |
| **TabICL** |  |  |  |  |
| HepG2 | 25% | 0.409±0.011 | 0.393±0.016 | 0.458 |
| HepG2 | 50% | 0.436±0.011 | 0.430±0.010 | 0.458 |
| HepG2 | 75% | 0.448±0.011 | 0.450±0.011 | 0.458 |
| Jurkat | 25% | 0.402±0.013 | 0.445±0.012 | 0.459 |
| Jurkat | 50% | 0.439±0.012 | 0.465±0.007 | 0.459 |
| Jurkat | 75% | 0.454±0.015 | 0.463±0.013 | 0.459 |
| RPE1 Essential | 25% | 0.503±0.016 | 0.630±0.020 | 0.583 |
| RPE1 Essential | 50% | 0.542±0.009 | 0.635±0.011 | 0.583 |
| RPE1 Essential | 75% | 0.568±0.011 | 0.607±0.013 | 0.583 |
| **TabPFN** |  |  |  |  |
| HepG2 | 25% | 0.408±0.009 | 0.404±0.019 | 0.456 |
| HepG2 | 50% | 0.438±0.012 | 0.442±0.010 | 0.456 |
| HepG2 | 75% | 0.445±0.011 | 0.455±0.013 | 0.456 |
| Jurkat | 25% | 0.388±0.013 | 0.439±0.007 | 0.451 |
| Jurkat | 50% | 0.426±0.012 | 0.456±0.010 | 0.451 |
| Jurkat | 75% | 0.445±0.015 | 0.453±0.015 | 0.451 |
| RPE1 Essential | 25% | 0.416±0.022 | 0.611±0.024 | 0.570 |
| RPE1 Essential | 50% | 0.496±0.002 | 0.612±0.011 | 0.570 |
| RPE1 Essential | 75% | 0.542±0.007 | 0.595±0.011 | 0.570 |

## References

A. K. Adduri, D. Gautam, B. Bevilacqua, A. Imran, R. Shah, M. Naghipourfar, N. Teyssier, R. Ilango, S. Nagaraj, M. Dong, et al. Predicting cellular responses to perturbation across diverse contexts with STATE. bioRxiv, pages 2025–06, 2025.

C. Ahlmann-Eltze, W. Huber, and S. Anders. Deep-learning-based gene perturbation effect prediction does not yet outperform simple linear baselines. Nature Methods, 22(8):1657–1661, Aug. 2025. ISSN 1548-7105. doi: 10.1038/s41592-025-02772-6. URL 10.1038/s41592-025-02772-6.

B. Anchang, A. Benz, D. Burkhardt, R. Cannoodt, M. Cortes, J. Fong, S. Kuppasani, R. Lieberman, T. Liu, M. Luecken, J. Mas-Rosario, R. Meinl, J. Nourisa, A. Szałata, F. Theis, J. Tumiel, T. Tunjic, M. Wang, N. Weber, and H. Zhao. A benchmark for prediction of transcriptomic responses to chemical perturbations across cell types. In Advances in Neural Information Processing Systems 37, NeurIPS 2024, page 20566–20616. Neural Information Processing Systems Foundation, Inc. (NeurIPS), 2024. doi: 10.52202/079017-0650. URL 10.52202/079017-0650.

M. Bühler, L. Purucker, and F. Hutter. Causal data augmentation for robust fine-tuning of tabular foundation models. arXiv [cs.LG], Jan. 2026.

C. Bunne, S. G. Stark, G. Gut, J. S. del Castillo, M. Levesque, K.-V. Lehmann, L. Pelkmans, A. Krause, and G. Rätsch. Learning single-cell perturbation responses using neural optimal transport. Nature Methods, 20(11):1759–1768, Sept. 2023. ISSN 1548-7105. doi: 10.1038/s41592-023-01969-x. URL 10.1038/s41592-023-01969-x.

E. Cole, G.-J. Huizing, S. Addagudi, N. Ho, E. Hasanaj, M. Kuijs, T. Johnstone, M. Carilli, A. Davi, C. Ellington, C. Feinauer, P. Li, R. Menegaux, S. Mohammadi, Y. Shao, J. Zhang, E. Lundberg, L. Song, Z. Bar-Joseph, and E. P. Xing. Foundation models improve perturbation response prediction. bioRxiv, 2026. doi: 10.64898/2026.02.18.706454. URL https://www.biorxiv.org/content/early/2026/02/19/2026.02.18.706454.

H. Cui, C. Wang, H. Maan, K. Pang, F. Luo, N. Duan, and B. Wang. scGPT: toward building a foundation model for single-cell multi-omics using generative AI. Nature methods, 21(8): 1470–1480, 2024.

M. Dong, A. Adduri, D. Gautam, C. Carpenter, R. Shah, C. Ricci-Tam, Y. Kluger, D. P. Burke, and Y. H. Roohani. Stack: In-context learning of single-cell biology. bioRxiv, 2026. doi: 10.64898/2026.01.09.698608. URL https://www.biorxiv.org/content/early/2026/01/09/2026.01.09.698608.

B. Feuer, R. T. Schirrmeister, V. Cherepanova, C. Hegde, F. Hutter, M. Goldblum, N. Cohen, and C. White. TuneTables: Context optimization for scalable prior-data fitted networks. arXiv [cs.LG], Feb. 2024.

N. Hollmann, S. Müller, L. Purucker, A. Krishnakumar, M. Körfer, S. B. Hoo, R. T. Schirrmeister, and F. Hutter. Accurate predictions on small data with a tabular foundation model. Nature, 637(8045):319–326, Jan. 2025. ISSN 1476-4687. doi: 10.1038/s41586-024-08328-6. URL 10.1038/s41586-024-08328-6.

Y. Ji, A. Tejada-Lapuerta, N. A. Schmacke, Z. Zheng, X. Zhang, S. Khan, I. Rothenaigner, J. Tschuck, K. Hadian, V. Hornung, and F. J. Theis. Scalable and universal prediction of cellular phenotypes enables in silico experiments. bioRxiv, 2025. doi: 10.1101/2024.08.12.607533. URL https://www.biorxiv.org/content/early/2025/09/23/2024.08.12.607533.

J.-H. Kim, C. S. Gibbs, S. Yun, H. O. Song, and K. Cho. Large-scale targeted cause discovery via learning from simulated data, 2024. URL https://arxiv.org/abs/2408.16218.

D. Klein, J. S. Fleck, D. Bobrovskiy, L. Zimmermann, S. Becker, A. Palma, L. Dony, A. Tejada-Lapuerta, G. Huguet, H.-C. Lin, N. Azbukina, F. Sanchís-Calleja, T. Uscidda, A. Szalata, M. Gander, A. Regev, B. Treutlein, J. G. Camp, and F. J. Theis. CellFlow enables generative single-cell phenotype modeling with flow matching. Apr. 2025.

J. Landsgesell, P. Knoll, and T. Wenzel. Scoringbench: A benchmark for evaluating tabular foundation models with proper scoring rules, 2026. URL https://arxiv.org/abs/2603.29928.

R. Littman, J. Levine, S. Maleki, Y. Lee, V. Ermakov, L. Qiu, A. Wu, K. Huang, R. Lopez, G. Scalia, T. Biancalani, D. Richmond, A. Regev, and J.-C. Hütter. Gene-embedding-based prediction and functional evaluation of perturbation expression responses with presage. 2025. doi: 10.1101/2025.06.03.657653. URL 10.1101/2025.06.03.657653.

M. Lotfollahi, A. Klimovskaia Susmelj, C. De Donno, L. Hetzel, Y. Ji, I. L. Ibarra, S. R. Srivatsan, M. Naghipourfar, R. M. Daza, B. Martin, J. Shendure, J. L. McFaline-Figueroa, P. Boyeau, F. A. Wolf, N. Yakubova, S. Günnemann, C. Trapnell, D. Lopez-Paz, and F. J. Theis. Predicting cellular responses to complex perturbations in high-throughput screens. Molecular Systems Biology, 19(6): e11517, 2023. doi: 10.15252/msb.202211517. URL https://www.embopress.org/doi/abs/10.15252/msb.202211517.

S. Müller, N. Hollmann, S. P. Arango, J. Grabocka, and F. Hutter. Transformers can do bayesian inference, 2024. URL https://arxiv.org/abs/2112.10510.

S. Müller, A. Reuter, N. Hollmann, D. Rügamer, and F. Hutter. Position: The future of bayesian prediction is prior-fitted, 2025. URL https://arxiv.org/abs/2505.23947.

A. Nadig, J. M. Replogle, A. N. Pogson, M. Murthy, S. A. McCarroll, J. S. Weissman, E. B. Robinson, and L. J. O’Connor. Transcriptome-wide analysis of differential expression in perturbation atlases. Nature Genetics, pages 1–10, 2025.

E. C. Neto. Using maximal information auxiliary variables to improve synthetic data generation based on tabPFN foundation models: preliminary results. In NeurIPS 2025 Workshop on Structured Probabilistic Inference & Generative Modeling, 2025. URL https://openreview.net/forum?id=sx0IcbKJk2.

G. Palla, S. Babu, P. Dibaeinia, J. D. Pearce, D. Li, A. A. Khan, T. Karaletsos, and J. M. Tomczak. Scalable single-cell gene expression generation with latent diffusion models, 2025. URL https://arxiv.org/abs/2511.02986.

J. Qu, D. Holzmüller, G. Varoquaux, and M. L. Morvan. TabICLv2: A better, faster, scalable, and open tabular foundation model. arXiv [cs.LG], Feb. 2026.

J. M. Replogle, R. A. Saunders, A. N. Pogson, J. A. Hussmann, A. Lenail, A. Guna, L. Mascibroda, E. J. Wagner, K. Adelman, G. Lithwick-Yanai, N. Iremadze, F. Oberstrass, D. Lipson, J. L. Bonnar, M. Jost, T. M. Norman, and J. S. Weissman. Mapping information-rich genotype-phenotype landscapes with genome-scale perturb-seq. Cell, 185(14):2559–2575.e28, 2022. ISSN 0092-8674. doi: 10.1016/j.cell.2022.05.013. URL 10.1016/j.cell.2022.05.013.

L. M. Saunders, S. R. Srivatsan, M. Duran, M. W. Dorrity, B. Ewing, T. H. Linbo, J. Shendure, D. W. Raible, C. B. Moens, D. Kimelman, and C. Trapnell. Embryo-scale reverse genetics at single-cell resolution. Nature, 623(7988):782–791, Nov. 2023.

M. Sextro, W. Kłos, and G. Dernbach. Mappfn: Learning causal perturbation maps in context, 2026. URL https://arxiv.org/abs/2601.21092.

O. Swelam, L. Purucker, J. Robertson, H. Raum, J. Boedecker, and F. Hutter. Does TabPFN understand causal structures? arXiv [cs.LG], Nov. 2025.

C. V. Theodoris, L. Xiao, A. Chopra, M. D. Chaffin, Z. R. Al Sayed, M. C. Hill, H. Mantineo, E. M. Brydon, Z. Zeng, X. S. Liu, et al. Transfer learning enables predictions in network biology. Nature, 618(7965):616–624, 2023.

G. Wang, T. Liu, J. Zhao, Y. Cheng, and H. Zhao. Modeling and predicting single-cell multi-gene perturbation responses with sclambda. Dec. 2024. doi: 10.1101/2024.12.04.626878. URL 10.1101/2024.12.04.626878.

H.-J. Ye, S.-Y. Liu, and W.-L. Chao. A closer look at tabpfn v2: Understanding its strengths and extending its capabilities, 2025. URL https://arxiv.org/abs/2502.17361.

J. Zhang, A. A. Ubas, R. de Borja, V. Svensson, N. Thomas, N. Thakar, I. Lai, A. Winters, U. Khan, M. G. Jones, et al. Tahoe-100m: A giga-scale single-cell perturbation atlas for context-dependent gene function and cellular modeling. BioRxiv, pages 2025–02, 2025.

R. Zhu, E. Dann, J. Yan, J. R. Retana, R. Goto, R. C. Guitche, L. K. Petersen, M. Ota, J. K. Pritchard, and A. Marson. Genome-scale perturb-seq in primary human cd4+ t cells maps context-specific regulators of t cell programs and human immune traits. bioRxiv, 2025. doi: 10.64898/2025.12.23.696273. URL https://www.biorxiv.org/content/early/2025/12/24/2025.12.23.696273.

